# Network Pharmacology and Bioinformatics Methods Reveal the Mechanism of Sishenwan in the Treatment of diabetic nephropathy

**DOI:** 10.1101/2023.02.14.528457

**Authors:** Yuanyuan Deng, Yu Ma, Sai Zhang, Mianzhi Zhang

**Author notes:** These authors contributed equally to this work.

## Abstract

*Objective*. In order to decipher the bioactive components and potential mechanisms of the traditional Chinese medicine (TCM) formula Sishenwan (SSW) for diabetic nephropathy (DN), we integrated network pharmacology and bioinformatics. *Methods*. The candidate compounds of SSW and relative targets were obtained from the TCMSP, BATMAN-TCM, SiwssTartgetPrediction, STITCH, and ChEMBL web servers. The UniProt database was used to translate the target names into gene names, and then constructed the herbal-compound-target network. DN-related targets were ascertained based on OMIM, CTD, GeneCards, DisGeNET, and GEO. Furthermore, there was a protein-protein interaction (PPI) network to explore the overlapping targets between SSW and DN, which focused on screening the pivotal targets by topology. GO and KEGG enrichment analyses were carried out to further understand the potential functions associated with the effect of SSW against DN. Eventually, molecular docking simulations were performed to validate the binding affinity between major bioactive components and hub genes. *Results.* A total of 120 candidate active compounds and 542 corresponding drug targets were derived, in which 195 targets intersected with DN. Then, KEGG pathway analysis showed that several signaling pathways were closely related to the process of SSW against DN, including the AGE-RAGE signaling pathway in diabetic complications, the TNF signaling pathway, and IL-17 signaling pathway, ect. The PPI network analysis identified *PTGS2, CREB1, ESR1, TNF, IL1B, INS, AKT1, PPARG*, and *JUN* were the top 9 hub targets. The molecular docking confirmed that the bioactive compounds of SSW had a firm binding affinity with hub targets. *Conclusions*. As a whole, the present study revealed that SSW exerted therapeutic effects on DN via modulating multi-targets with multi-compounds through multi-pathways.

## 1. Introduction

As one of the most important microvascular complications of diabetes, diabetic nephropathy (DN) can cause the continuous decline of renal functions and irreversible damage to pathological structures^[1]^. The control of blood sugar, blood pressure, and blood lipids is the basic treatment of DN and has been widely applied in clinical practice^[2]^. However, these therapeutic means cannot effectively prevent the end-stage progression of DN, and there are limited treatment options for patients with DN. Therefore, it is urgently needed to develop new clinical intervention measures to delay the progression of renal functions.

As is known to all, the clinical application of traditional Chinese medicine (TCM) dates back thousands of years. TCM is featured by good clinical efficacy, few side effects and low drug resistance, which continuously expands the application range of Chinese patent medicine and makes TCM increasingly recognized all over the world^[3]^. It is extensively used in the prevention and treatment of DN in China. TCM is good at combining a holistic view with treatment based on syndrome differentiation and has advantages of multi-components, multi-targets, and multi-approaches in the treatment of internal medicine diseases^[4]^. In recent years, many scholars in the world have been actively exploring the potential mechanism of TCM compound therapy for DN. Some convincing preclinical and clinical evidence shows that TCM has extensive pharmacological action and good curative effect in the treatment of DN^[3–5]^. This indicates that TCM compound therapy can get a good curative effect on DN. SSW was first recorded in *Chen’s Prescription for Infantile Pox and Eruption by Chen Wenzhong in the Southern Song Dynasty*^[6]^. According to the book, SSW is composed of fructus psoraleae, fructus evodiae, myristica fragrans, schisandra chinensis and Chinese date^[6]^, and it can tonify the kidney and spleen. Besides, the above-mentioned five Chinese herbal medicinal ingredient ingredients are commonly used in the treatment of DN in China^[5, 7–9]^. However, the complexity of TCM compounds and the diversity of their mechanisms of action limit their clinical application prospects.

Network pharmacology and bioinformatics is a biological network based on interdisciplinary fusion and data mining which can elucidate the complex synergistic effects between “drug-target-disease” from a deep, multi-angle, and all-round perspective^[10, 11]^. Therefore, network pharmacology and bioinformatics were used to systematically integrate drug and disease information. Besides, the molecular mechanism and efficacy of SSW in treating DN were deeply explored from a macro perspective. The binding mode and affinity of the complex were accurately simulated and predicted by combining with molecular docking technology to verify the results of this study and provide a reference for the follow-up research and development of new drugs and clinical treatment.

## 2. Materials and Methods

### 2.1. Screening of Active Ingredients and Target Genes

All biological active ingredients of SSW were identified by using TCMSP data platform^[12]^ (http://tcmspw.com/tcmsp.php) and BATMAN Website (http://bionet.ncpsb.org/batman-tcm/)^[13]^. The potential targets related to SSW were obtained from the following databases: 1) TCMP, 2) SiwssTartgetPrediction^[14]^ (http://swisstargetprediction.ch/), 3) STITCH^[15]^ (http://stitch.embl.de), and (4) the ChEMBL^[16]^ (https://www.ebi.ac.uk/chembl/). With the help of UniProt database^[17]^ (https://www.uniprot.org/), the above drug target proteins were converted into their corresponding standard genetic symbols (Species setting: “human species”). Moreover, the component-target network was constructed by using Cytoscape^[18, 19]^ (https://cytoscape.org/, version 3.8.2).

### 2.2. Collection of DN Targets

First of all, the disease targets were searched with the keyword of “Diabetic nephropathy” in the database of DisGeNET^[20]^ (https://www.disgenet.org), GeneCards^[21]^ (https://www.genecards.org), OMIM^[22]^ (https://www.omim.org) and CTD^[23]^ (http://ctd.mdibl.org/). In addition, the differentially expressed genes (DEGs) were identified through the Comprehensive Gene Expression database^[24]^ (http://www.ncbi.nlm.nih.gov/GEO/). The existing transcriptional RNA-seq data for DN were analyzed in the gene expression dataset GSE30528, GSE1009, GSE96804, and GSE47183 (Species setting: “human species”).The gene chip data was analyzed by using Sva and Limma of Rmur3.6.2^[25]^, which met the Screening criteria of log2(foldchange)|>1 and P<0.05. These genes were considered to be distinct DEGs. The DEGs were visualized by using the “ggpubr” and “ggthemes” in R language package. Next, all the DECs were integrated and sorted through RobustRankAggreg^[26]^ (https://cran.rstudio.com/bin/windows/contrib//3.5/RoburRankAggreg1.1.zip). The obtained genes were then integrated with the disease genes, and all the targets of DN were obtained. Compound Disease Targets of SSW in treating DN were got by intersection with the compound targets of SSW.

### 2.3. Protein-Protein Interaction (PPI) Network and Topological Analysis

To expound the interaction of target proteins in the whole network, compound targets were uploaded to the STRING11.0 database^[27]^ (https://string-db.org) to construct a PPI network. And the filtering criteria were set as Homo sapiens and confidence≥0.4. The PPI networks consisting of overlapping component targets were constructed in Cytoscape. The key topology parameters of nodes were evaluated through the network analyzer in Cytoscape. The nodes with high degree values indicated that they were considered critical targets in the PPI. Then, the composite target was analyzed by using MCODE plug-in and the Degree of protein association in the module was scored by using the following criteria: degree Cutoff≥2, Node ScoreCutoff=0.2, K-core≥2, and Max Depth=100.

### 2.4. Enrichment Analysis of Composite Targets

The enrichment analysis of composite targets was performed in the DAVID online database (https://David.ncifcrf.gov/summary.jsp). A value of *P* < 0.05 was considered significant. The three categories of biological process (BP), cellular component (CC), and molecular function (MF) were the core elements of GO functional analysis. The first 20 items were selected for visualization. The component-target-pathway network was drawn by Cytoscape.

### 2.5. Docking of Molecules

Based on the results of the component-target-pathway network, the molecular docking of our core target protein was conducted. The small molecular ligands (Mol2 format) of biological active ingredients in SSW were obtained from PubChem database^[28]^ (https://PubChem.ncbi.nlm.nih.gov/). Besides, the two-dimensional and three-dimensional structures of proteins were downloaded from the RCSB PDB database^[29]^ (http://www.RCSB.org/). The preprocessing of proteins was performed through Syby-X software, including dehydration and hydrogenation, and then saved as PDBQT files. After completing the above steps, they were visualized using PyMOL software (https://www.pymol.org).

## 3. Results

### 3.1. Screening of Active Ingredients and Targets in SSW

From the TCMSP and BATMAN-TCM platforms, a total of 120 effective bioactive compounds (including 9 fructus psoralea, 27 Chinese dates, 31 myristica fragrans, 53 schisandrum chinensis and 30 fructus evodiaes) were identified by SSW, which corresponded to 542 targets. The active ingredient-target network was constructed by using Cytoscape software. After removing isolated nodes, the network consisted of 320 nodes and 1197 edges, where blue polygons, green circles, and red polygons represented components, target genes, and herbs, respectively (Figure 2). The degree value of a node represented the number of nodes connected to nodes in the network. The higher the degree value, the more connections a node had. Additional information on each candidate bioactive ingredient is presented in Supplement Table 1.

**Figure 1:**
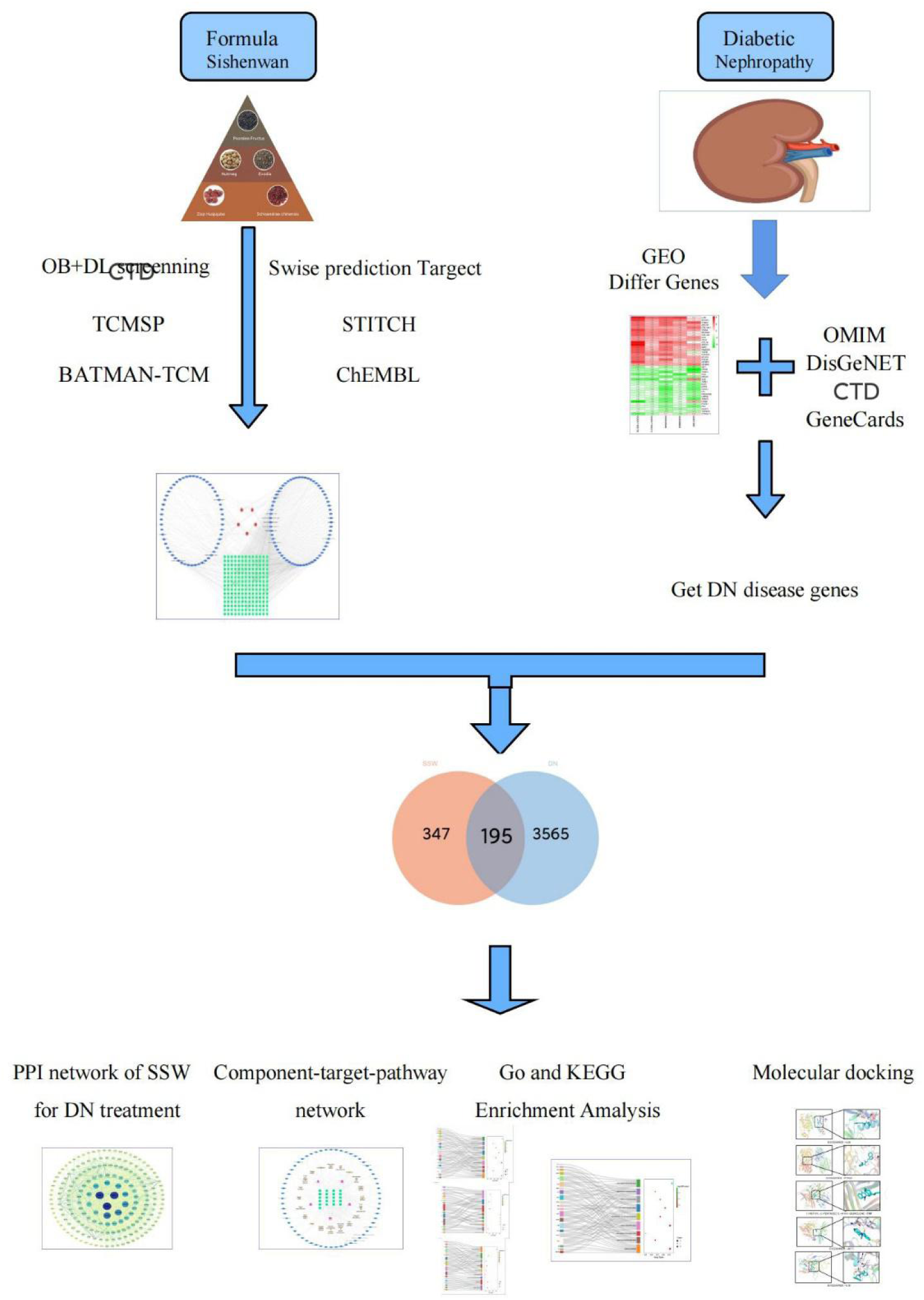
Network-based framework for pharmacology integration strategies.

**Figure 2:**
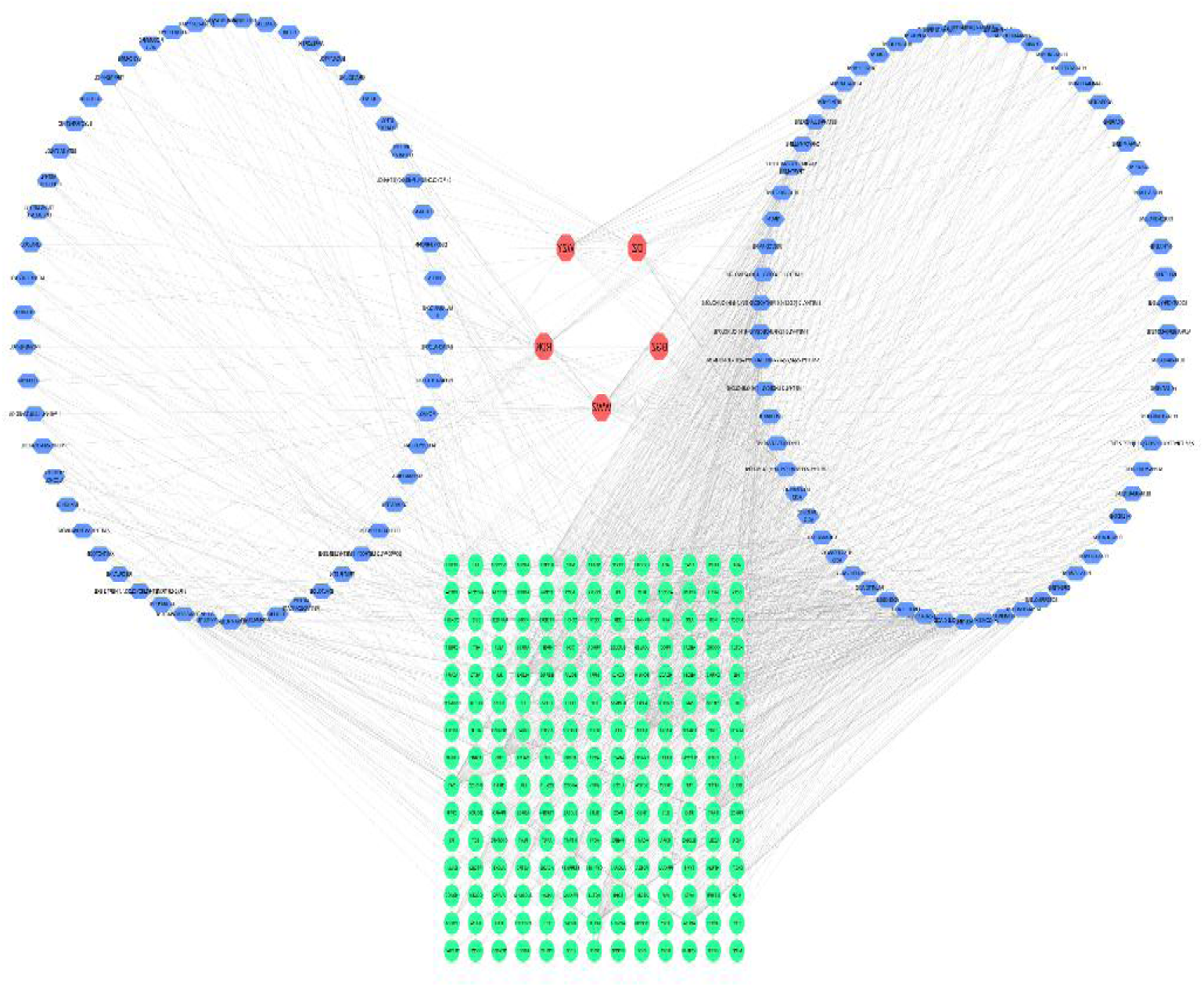
Compound-target network: red circles represent the herbs in SSW; hexagons represent active compounds of each herb; green circles represent related targets (The components are presented in Table 1).

**Table 1:**
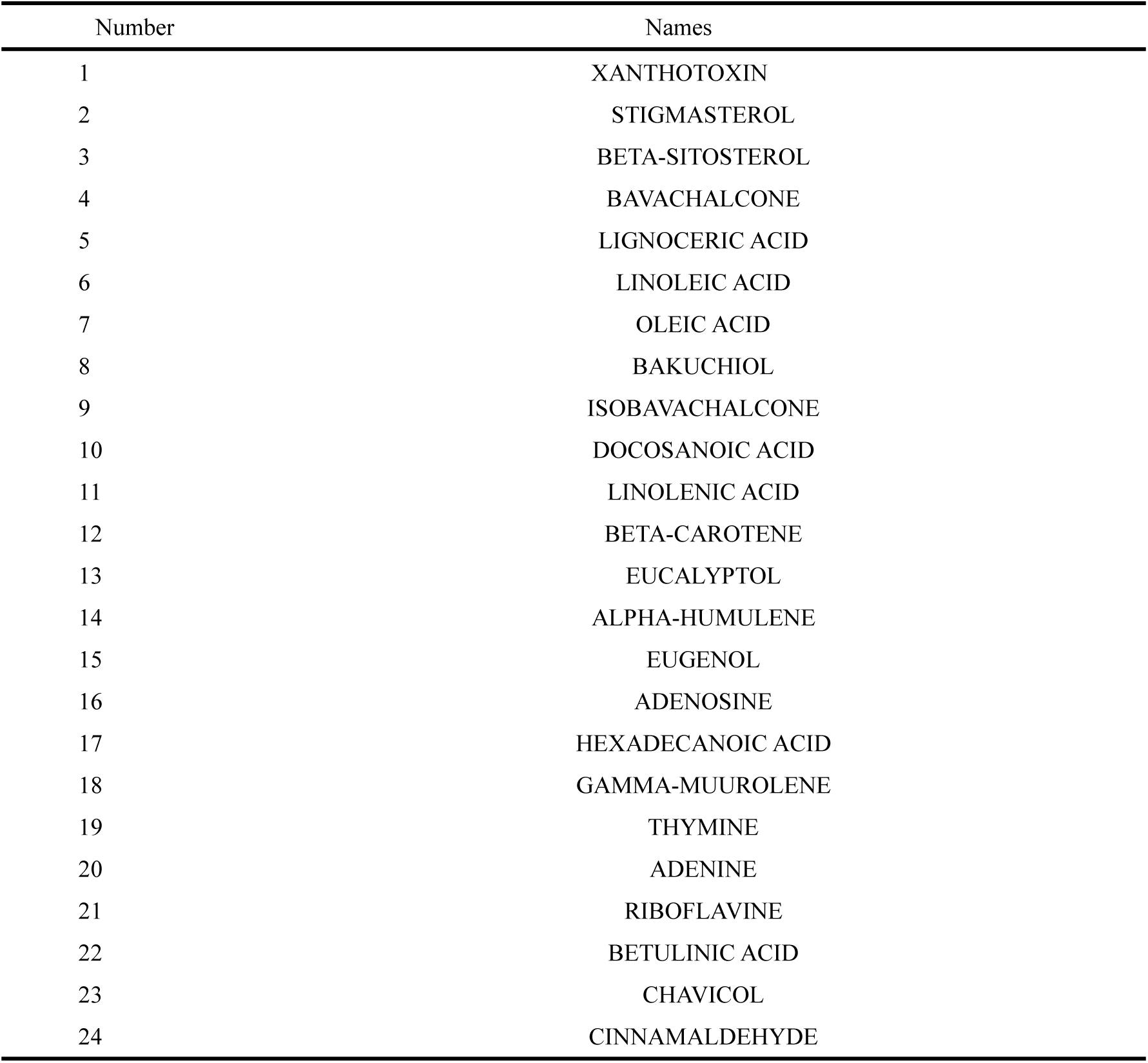
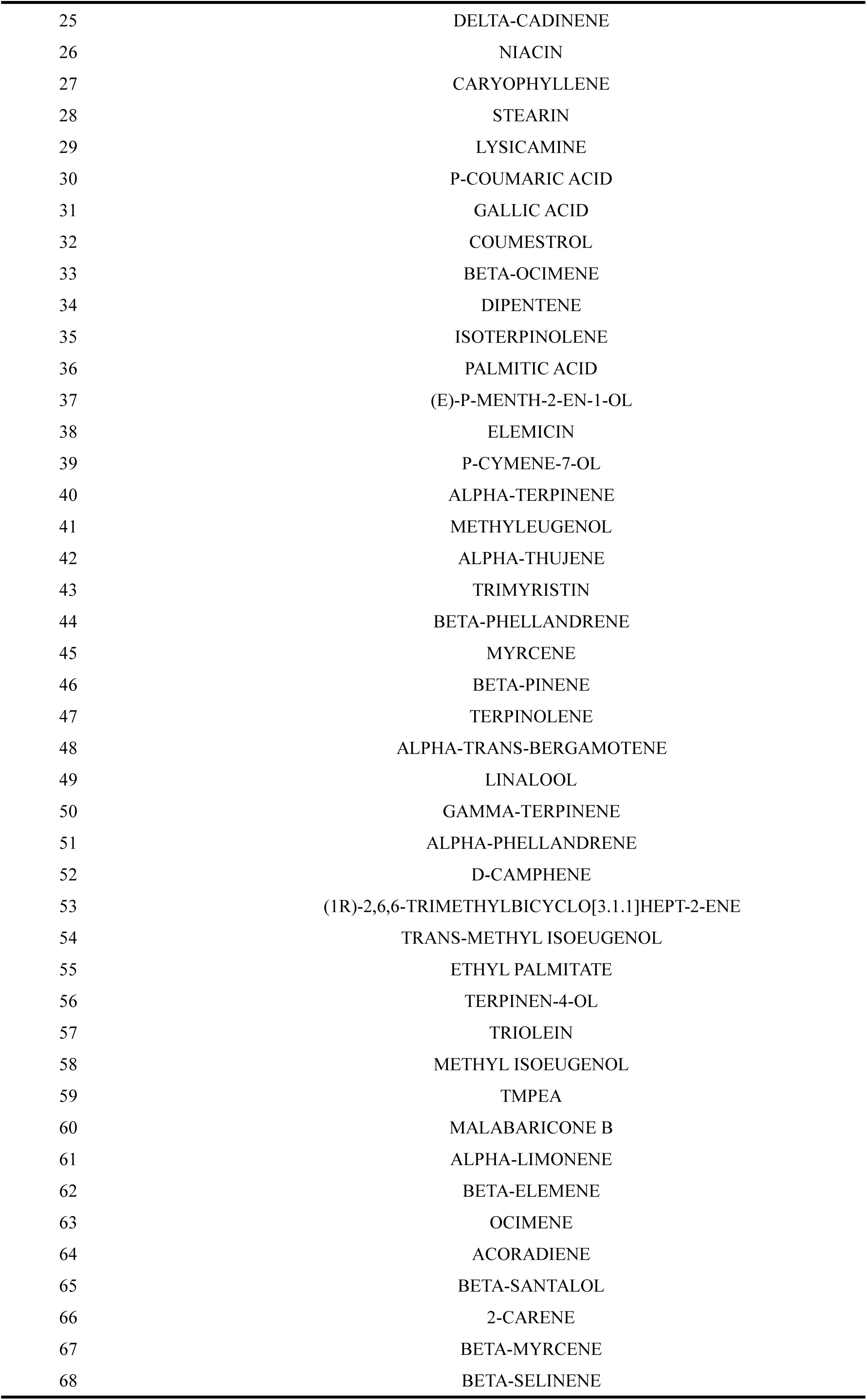
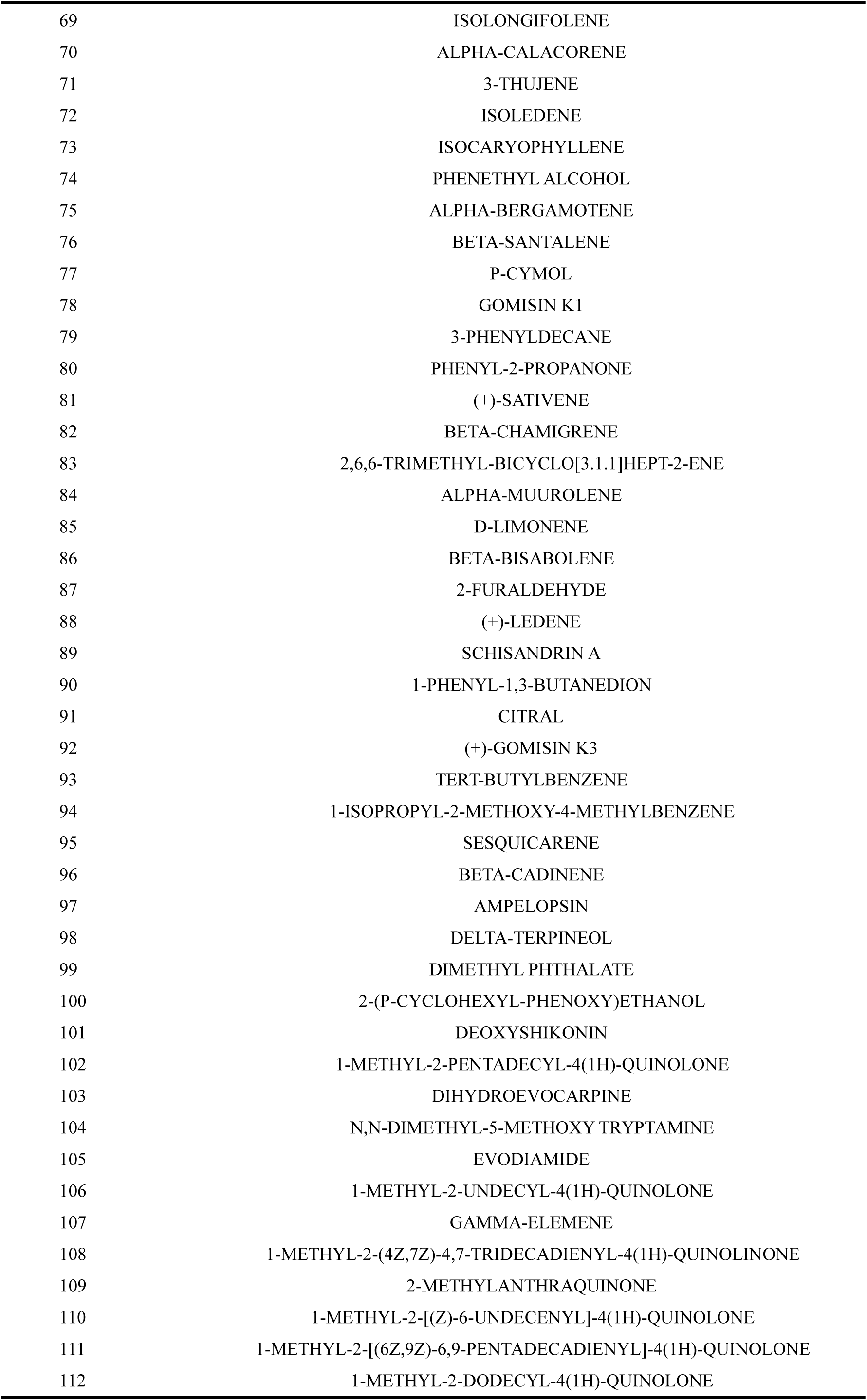
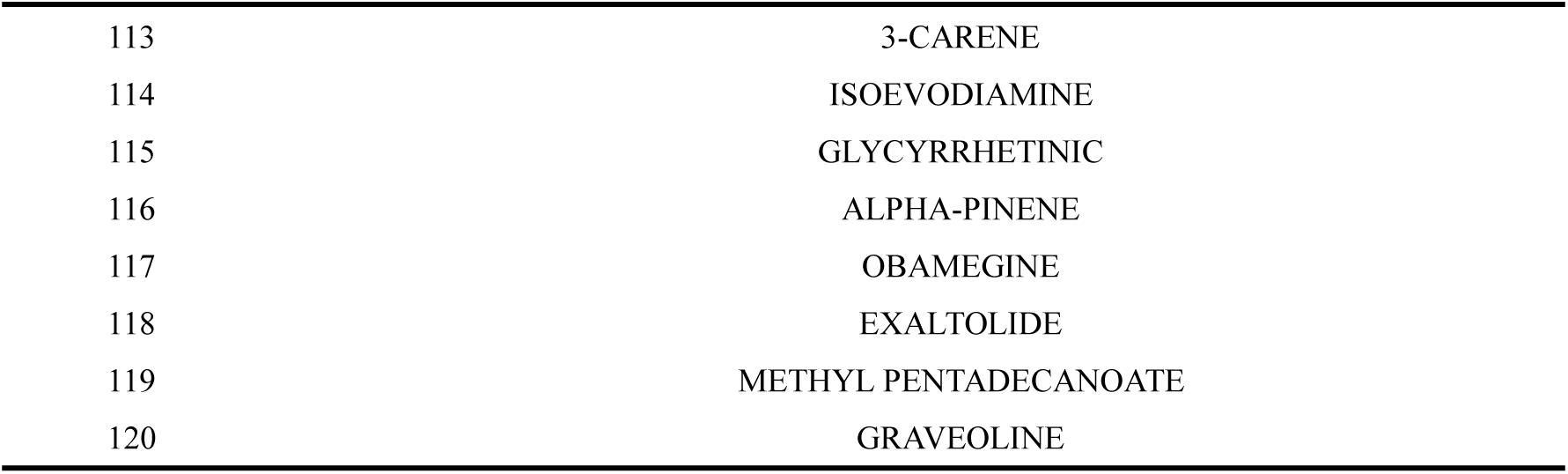
Bioactive compounds of SSW in DN therapy.

### 3.2. DN-associated targets and candidate genes

A total of 3236 DN targets were obtained from GeneCards database, and 3760 disease targets were got by supplementing DisGeNET, OMIM, and CTD numbers. The datasets GSE30528, GSE1009, GSE96804 and GSE47183 that met the screening criteria were downloaded from the GEO database. The RRA package was used for integrated analysis of the four datasets, and 82 robust DEGs were determined, among which 46 were up-regulated and 36 were down-regulated (Figure 3 and Supplement Table 2). When combined with the above disease targets, a total of 3760 DN targets were collected after removing duplicate values. Then, the predicted targets of DN were mapped to the targets of SSW, and 195 intersection targets were identified. Among them, 195 overlapping targets between SM and DN were identified and collected for further mechanism investigation (Figure 4 and Supplement Table 3).

**Figure 3:**
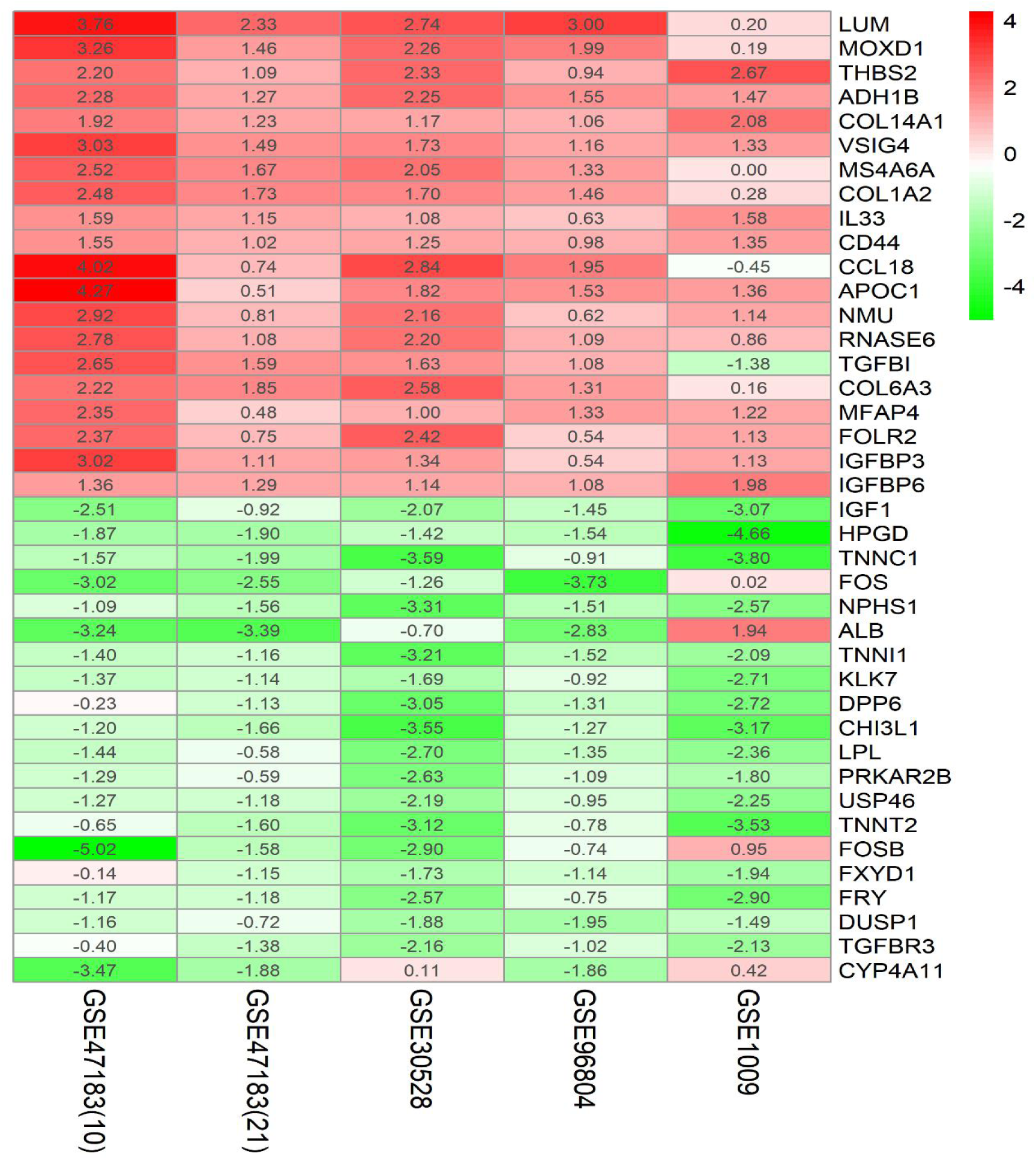
Heat maps of the first 20 up-regulated and down-regulated robust DEGs; Green represents down-regulated robust DEGs, while red represents up-regulated robust DEGs. The value in the box is the log2FC value of the robust DEGs.

**Figure 4:**
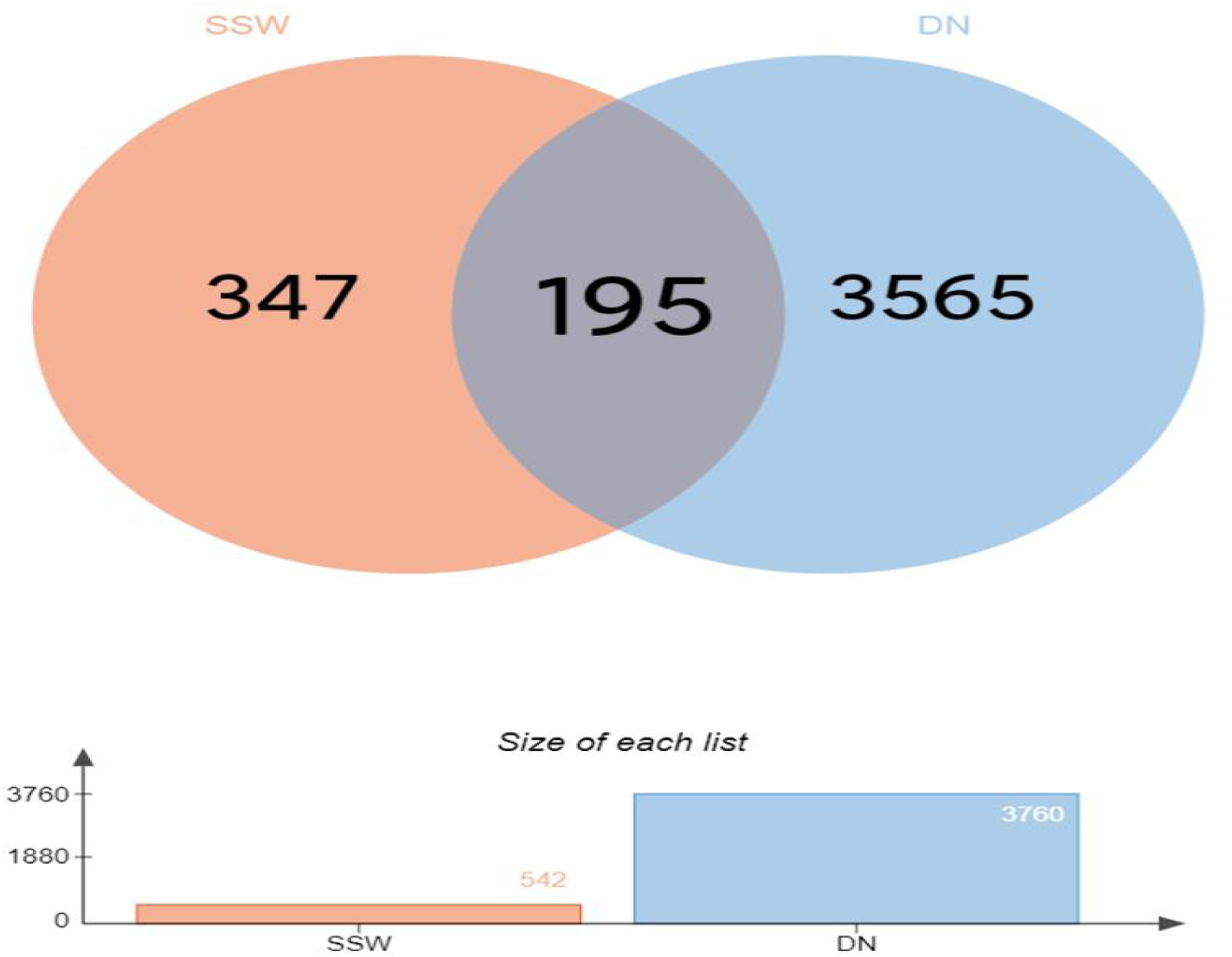
Venn’s diagram of intersection targets of SSW and DN.

### 3.3 PPI Network

The overlapping targets were imported into the STRING database. The PPI network (Figure 5 and Supplement Table 4) had 195 nodes and 1271 edges. Topology analysis was performed by using cytoHubba plug-in (https://apps.cytoscape.org/apps/cytohubba). The top 10 Hub genes, including INS, AKT1, TNF, IL1B, PPARG, JUN, CREB1, PTGS2, and ESR1 were regarded as key genes for SSW treatment. The darker the color, the higher the degree. Besides, cluster analysis was performed through MCODE plug-in (Table 2). MCODE plug-in screened out 5 modules and the core module cluster 1 with the highest Score was determined, which was composed of 24 nodes and 176 edges (Figure 5). The core target group of the active components in SSW (including TNF, AKT1, JUN, ESR1, PPARG, etc.) scored 15.304. The cluster score represented the core density of nodes and topologically adjacent nodes. The higher the score, the more concentrated the cluster, indicating that SSW might play a role through the multi-target synergistic mechanism with DN targets. The top 10 genes in degree and cluster1 genes were intersected to obtain the final core target genes: PTGS2, CREB1, ESR1, TNF, IL1B, INS, AKT1, PPARG, and JUN.

**Figure 5:**
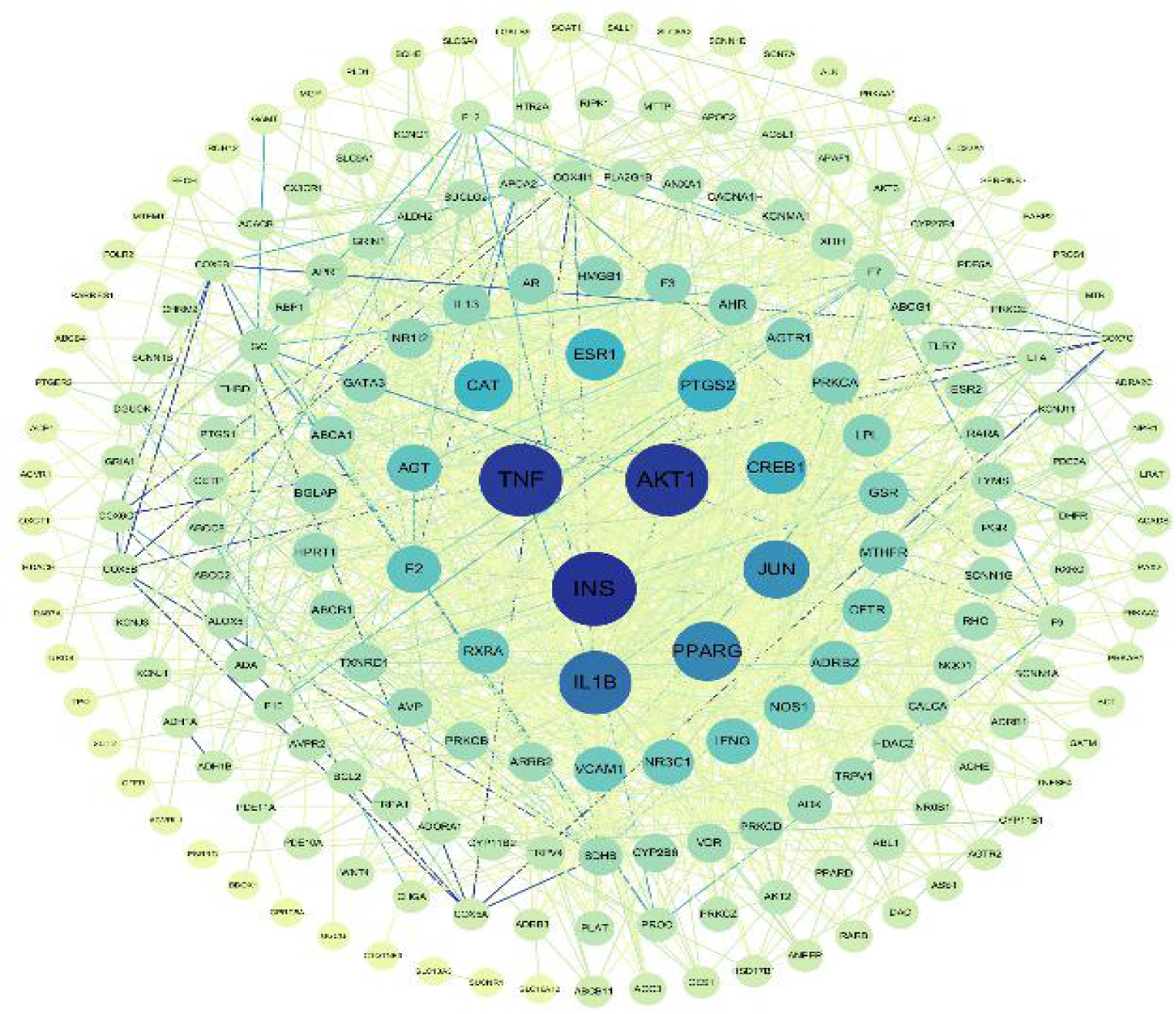
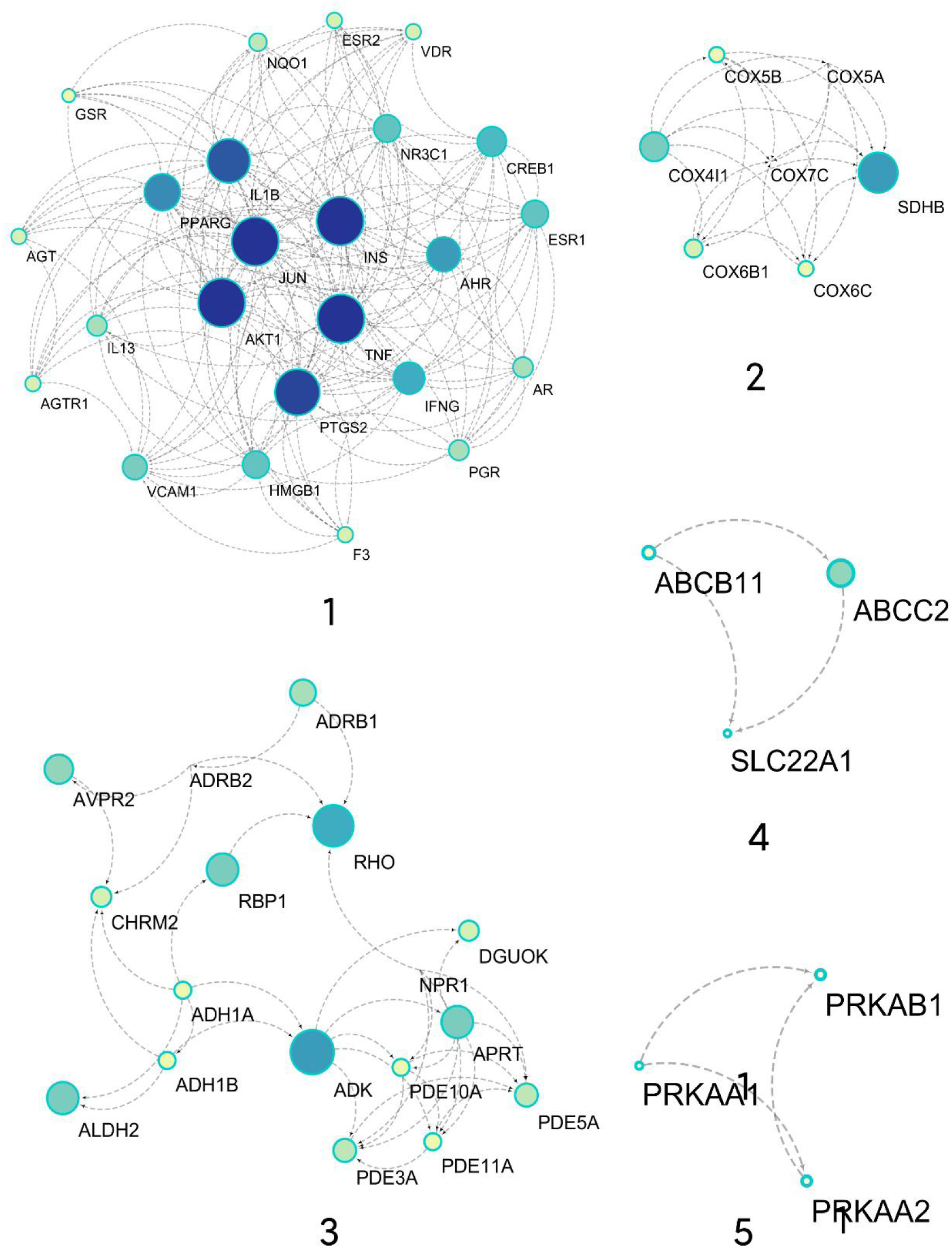
PPI network of SSW for DN treatment; Circles represent genes; lines represent the interactions between genes. The line colors represent the types of protein interactions; the area of the node is proportional to the degree value.

**Table 2:** Five gene clusters of SSW for DN.

| Cluster | Score | Nodes | Edges | Count |
| --- | --- | --- | --- | --- |
| Cluster 1 | 15.304 | 24 | 176 | 24 |
| cluster 2 | 7 | 7 | 21 | 7 |
| cluster 3 | 4.5 | 17 | 36 | 17 |
| cluster 4 | 3 | 3 | 3 | 3 |
| cluster 5 | 3 | 3 | 3 | 3 |

**Table 3:**
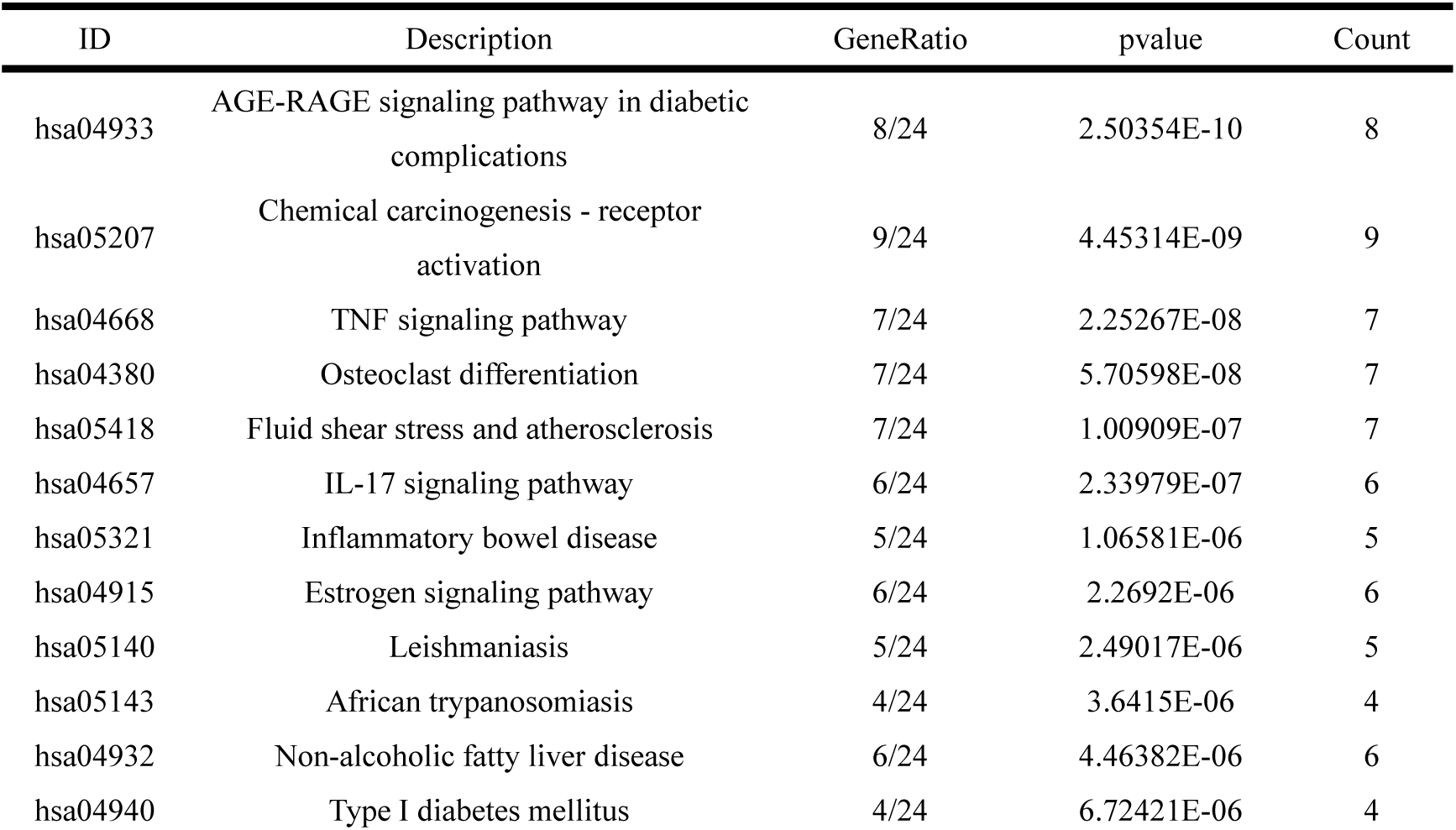
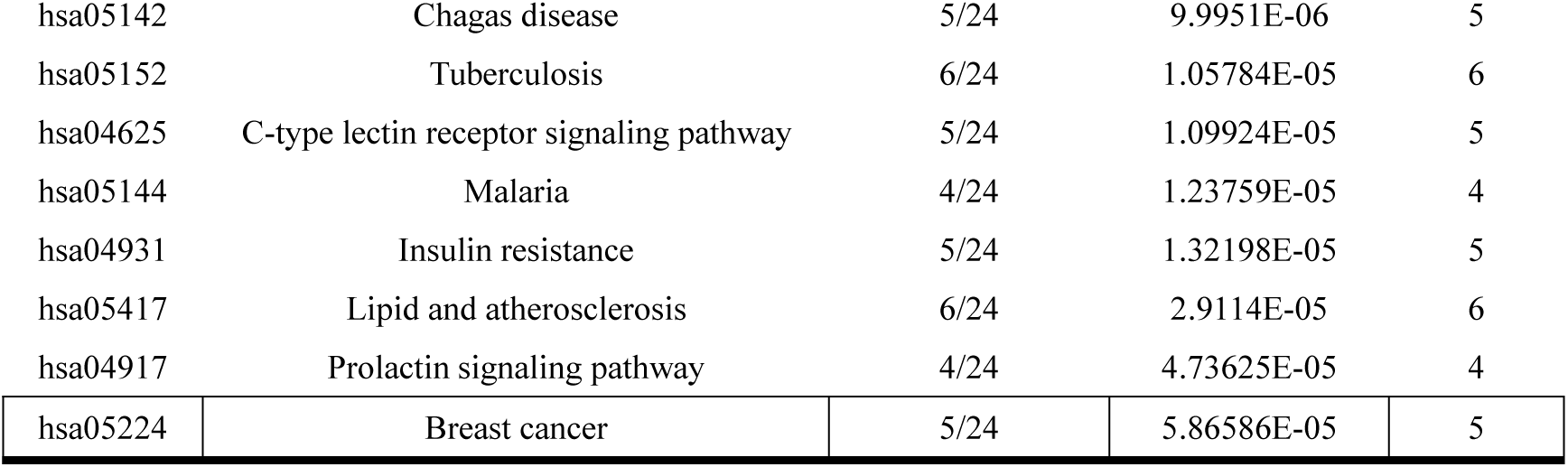
Top 20 pathways.

| ID | Description | GeneRatio | pvalue | Count |
| --- | --- | --- | --- | --- |
| hsa04933 | AGE-RAGE signaling pathway in diabetic complications | 8/24 | 2.50354E-10 | 8 |
| hsa05207 | Chemical carcinogenesis - receptor activation | 9/24 | 4.45314E-09 | 9 |
| hsa04668 | TNF signaling pathway | 7/24 | 2.25267E-08 | 7 |
| hsa04380 | Osteoclast differentiation | 7/24 | 5.70598E-08 | 7 |
| hsa05418 | Fluid shear stress and atherosclerosis | 7/24 | 1.00909E-07 | 7 |
| hsa04657 | IL-17 signaling pathway | 6/24 | 2.33979E-07 | 6 |
| hsa05321 | Inflammatory bowel disease | 5/24 | 1.06581E-06 | 5 |
| hsa04915 | Estrogen signaling pathway | 6/24 | 2.2692E-06 | 6 |
| hsa05140 | Leishmaniasis | 5/24 | 2.49017E-06 | 5 |
| hsa05143 | African trypanosomiasis | 4/24 | 3.6415E-06 | 4 |
| hsa04932 | Non-alcoholic fatty liver disease | 6/24 | 4.46382E-06 | 6 |
| hsa04940 | Type I diabetes mellitus | 4/24 | 6.72421E-06 | 4 |
| hsa05142 | Chagas disease | 5/24 | 9.9951E-06 | 5 |
| hsa05152 | Tuberculosis | 6/24 | 1.05784E-05 | 6 |
| hsa04625 | C-type lectin receptor signaling pathway | 5/24 | 1.09924E-05 | 5 |
| hsa05144 | Malaria | 4/24 | 1.23759E-05 | 4 |
| hsa04931 | Insulin resistance | 5/24 | 1.32198E-05 | 5 |
| hsa05417 | Lipid and atherosclerosis | 6/24 | 2.9114E-05 | 6 |
| hsa04917 | Prolactin signaling pathway | 4/24 | 4.73625E-05 | 4 |
| hsa05224 | Breast cancer | 5/24 | 5.86586E-05 | 5 |

### 3.4. Results of Enrichment Analysis

Based on GO enrichment analysis (Figure 6), 195 core targets were identified, involving molecular function (MF), cell component (CC), and biological processes (BP). To be specific, MF mainly involved nuclear receptor activity (GO:0004879), ligand-activated transcription factor activity (GO:0098531) and transcription coactivator binding (GO:0001223); BP mainly involved positive regulation of lipid metabolic process (GO:0045834), positive regulation of oxidoreductase activity (GO:0051353), regulation of oxidoreductase activity (GO:0051341), polymerase II transcription regulator complex (GO:0090575), and the external side of plasma membrane (GO:0009897). Moreover, the main signaling pathways involved in DN treatment were identified through KEGG enrichment analysis (Figure 7 and table 2). These included AGE-RAGE signaling pathway in diabetic complications (hsa04933), TNF signaling pathway (hsa04668), fluid shear stress and signaling pathway in AGE-RAGE atherosclerosis (hsa05418), and IL-17 signaling pathway (hsa04657) (Figure 7). AGE-RAGE signaling pathway in diabetic complications (hsa04933) was the most highly enriched and closely associated with other pathways and it was identified as the core pathway for SSW to function. AGE-RAGE signaling pathway in diabetic complications is shown in Figure 8. A detailed description of KEGG enrichment pathways is depicted in Supplement Table 5.

**Figure 6:**
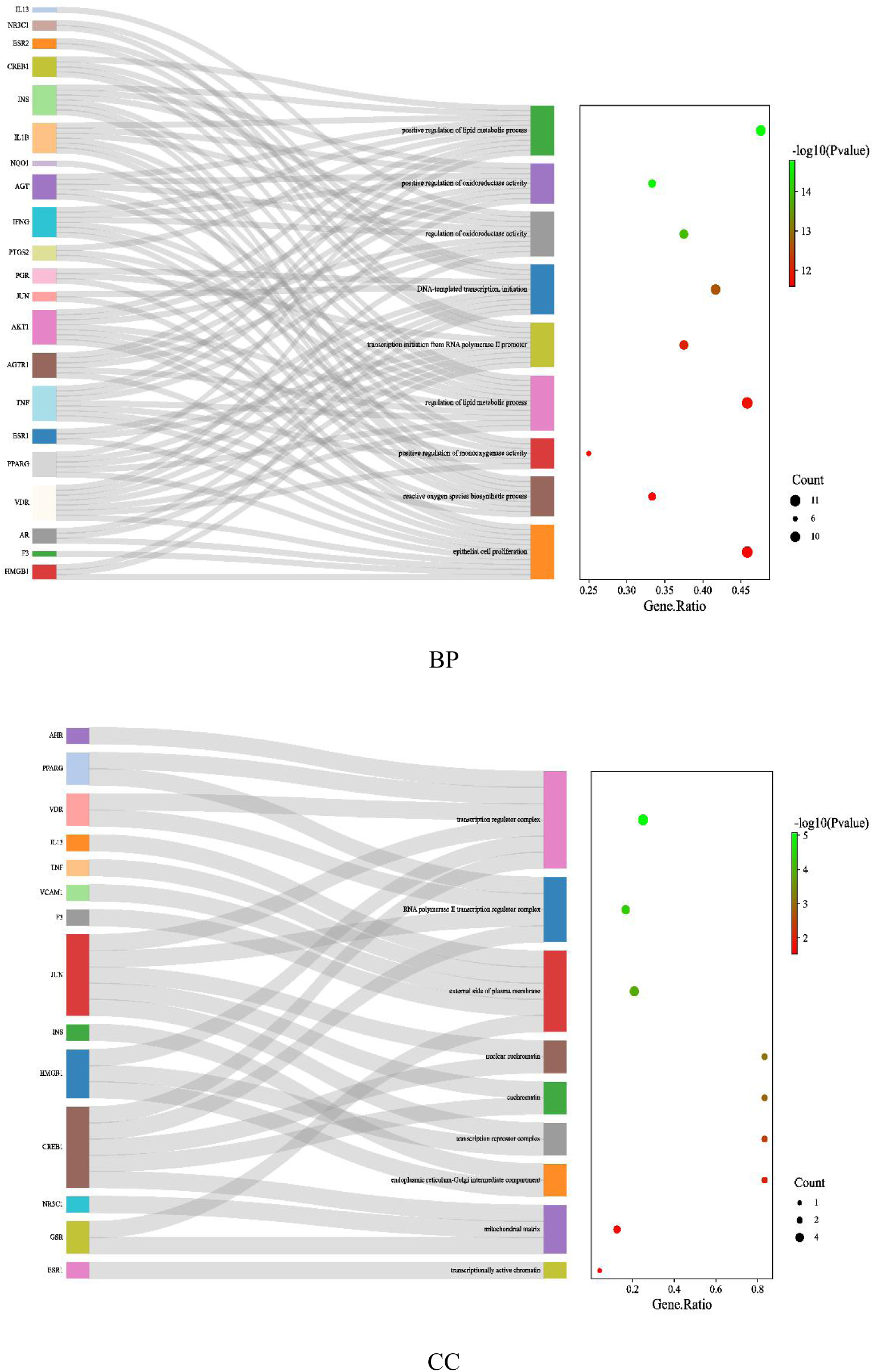
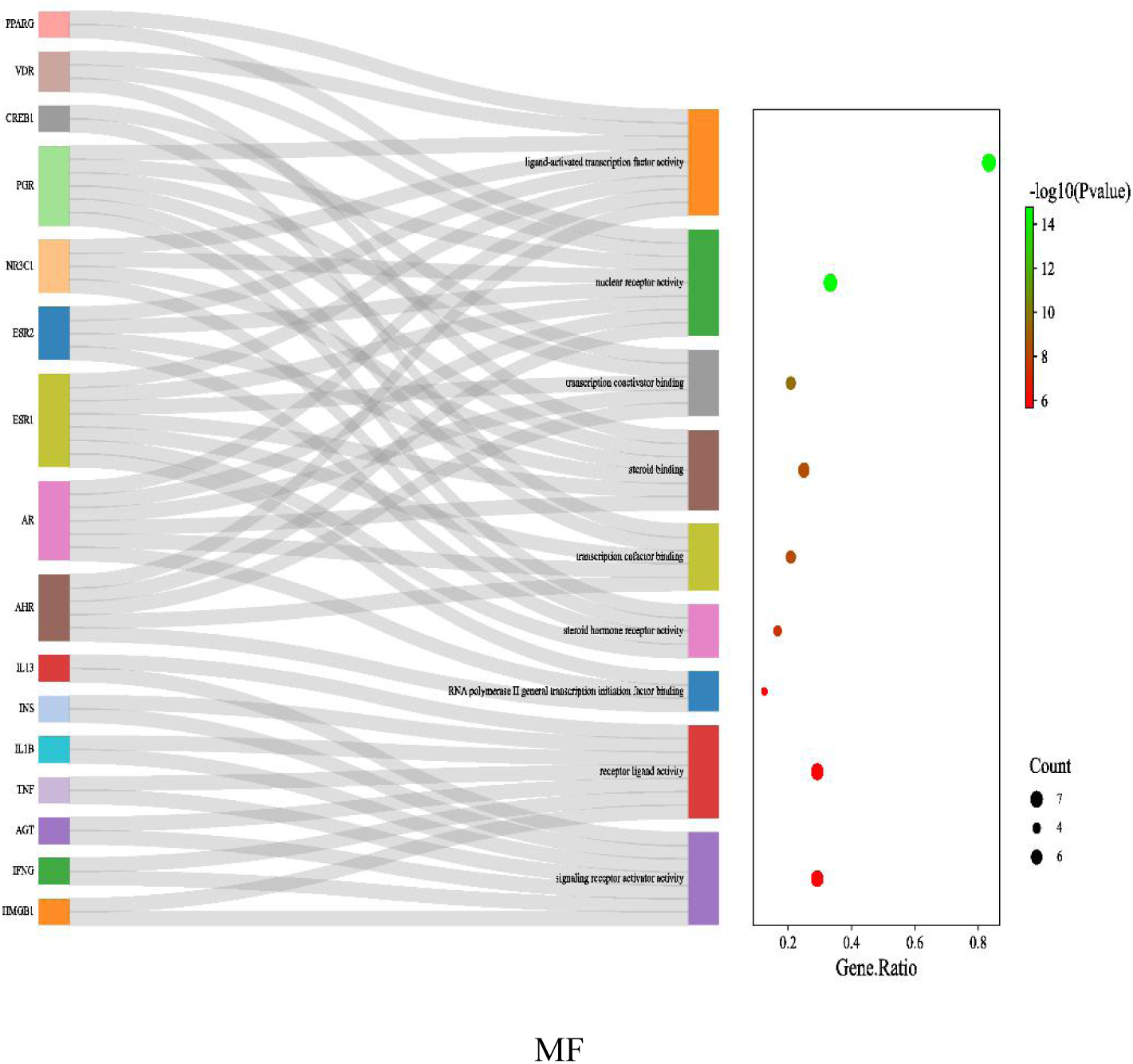
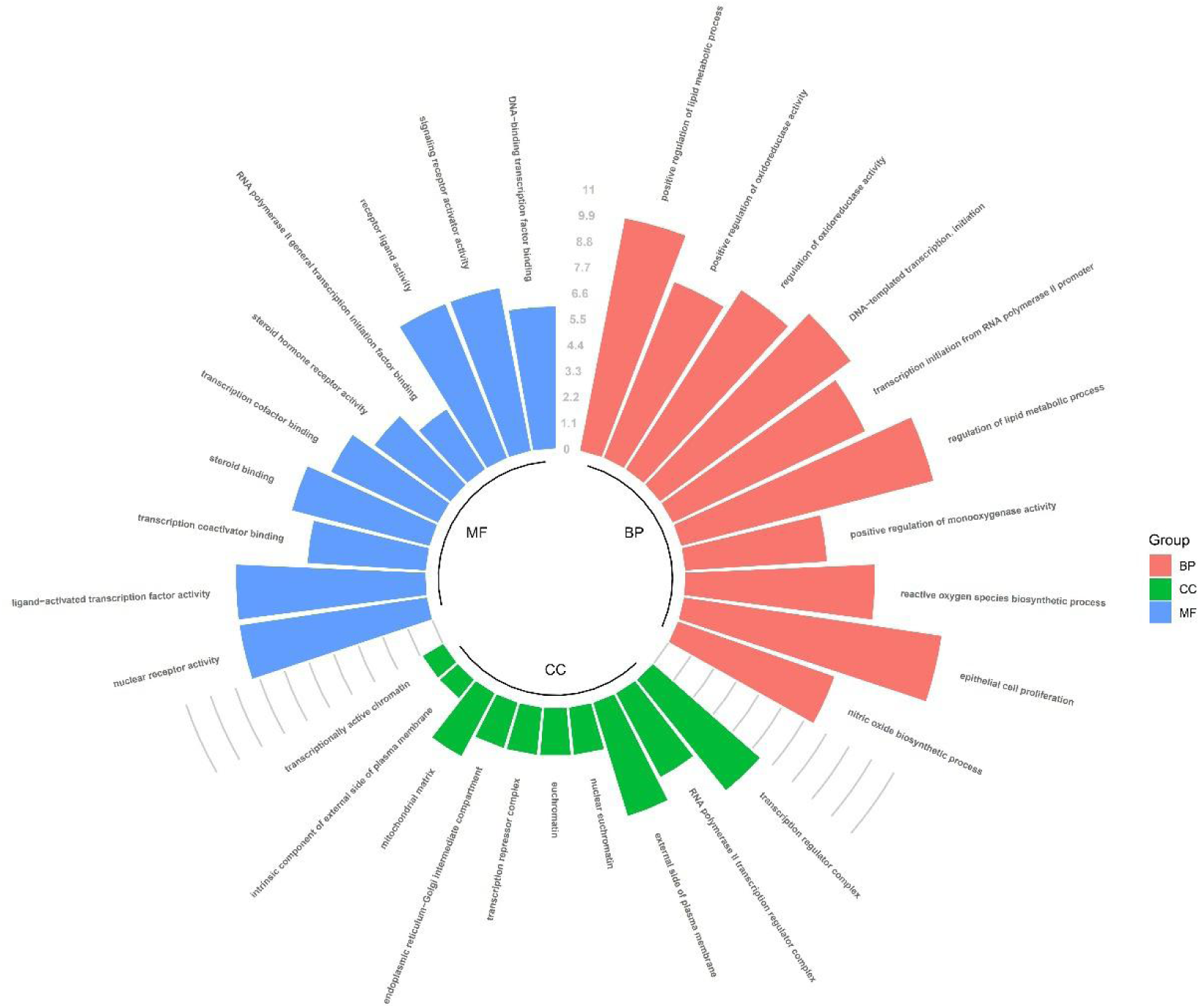
The 10 most significance of GO analysis.

**Figure 7:**
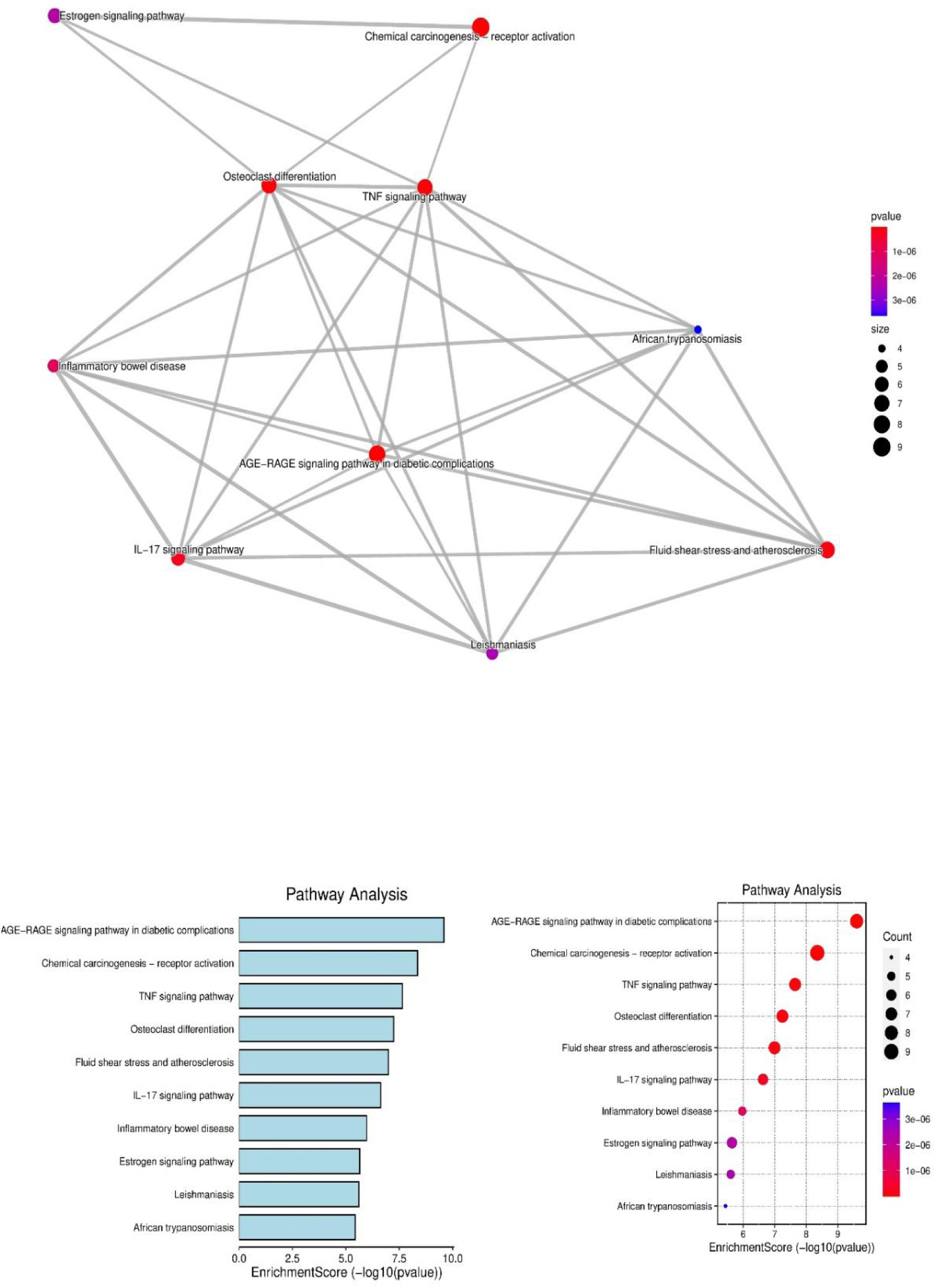
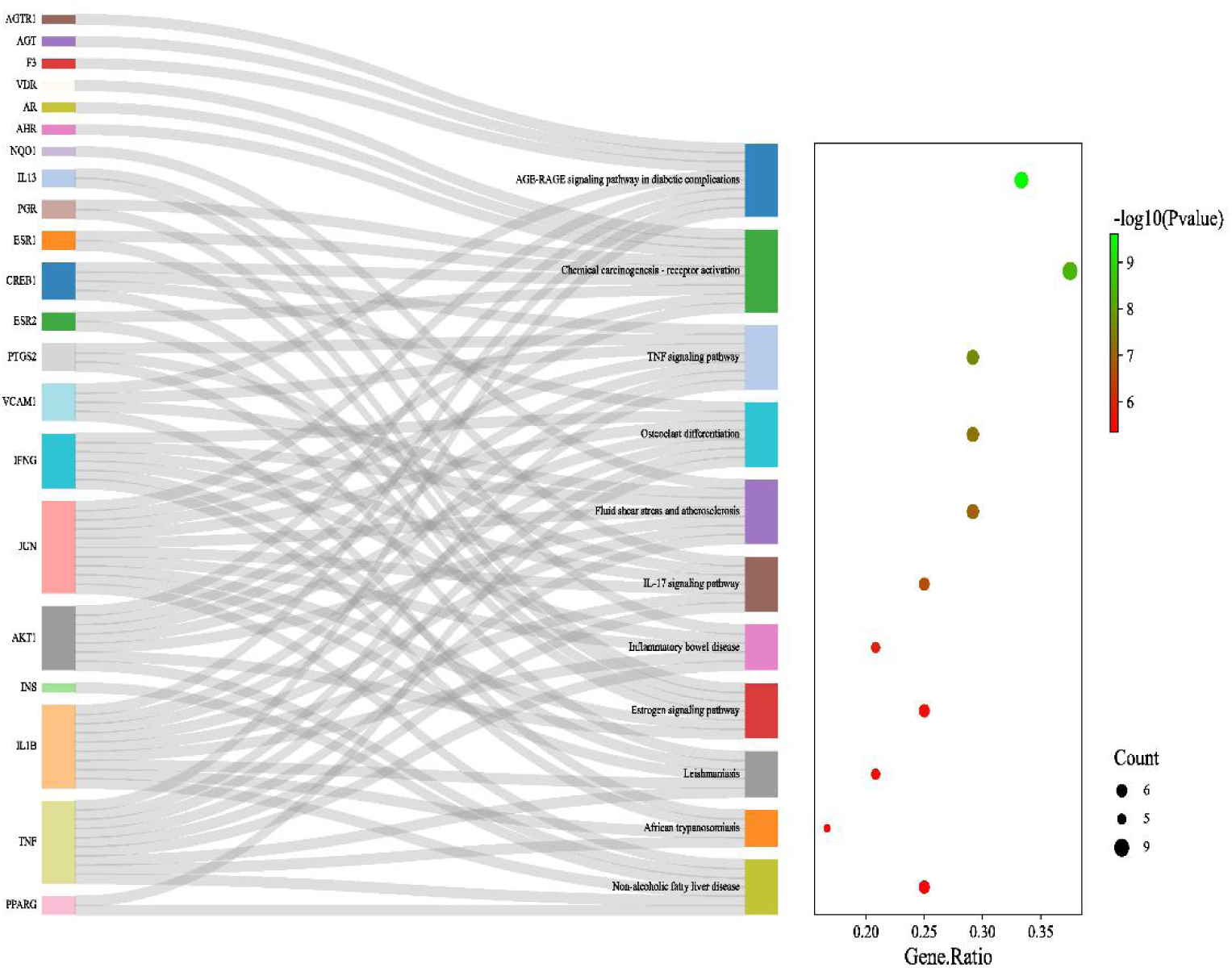
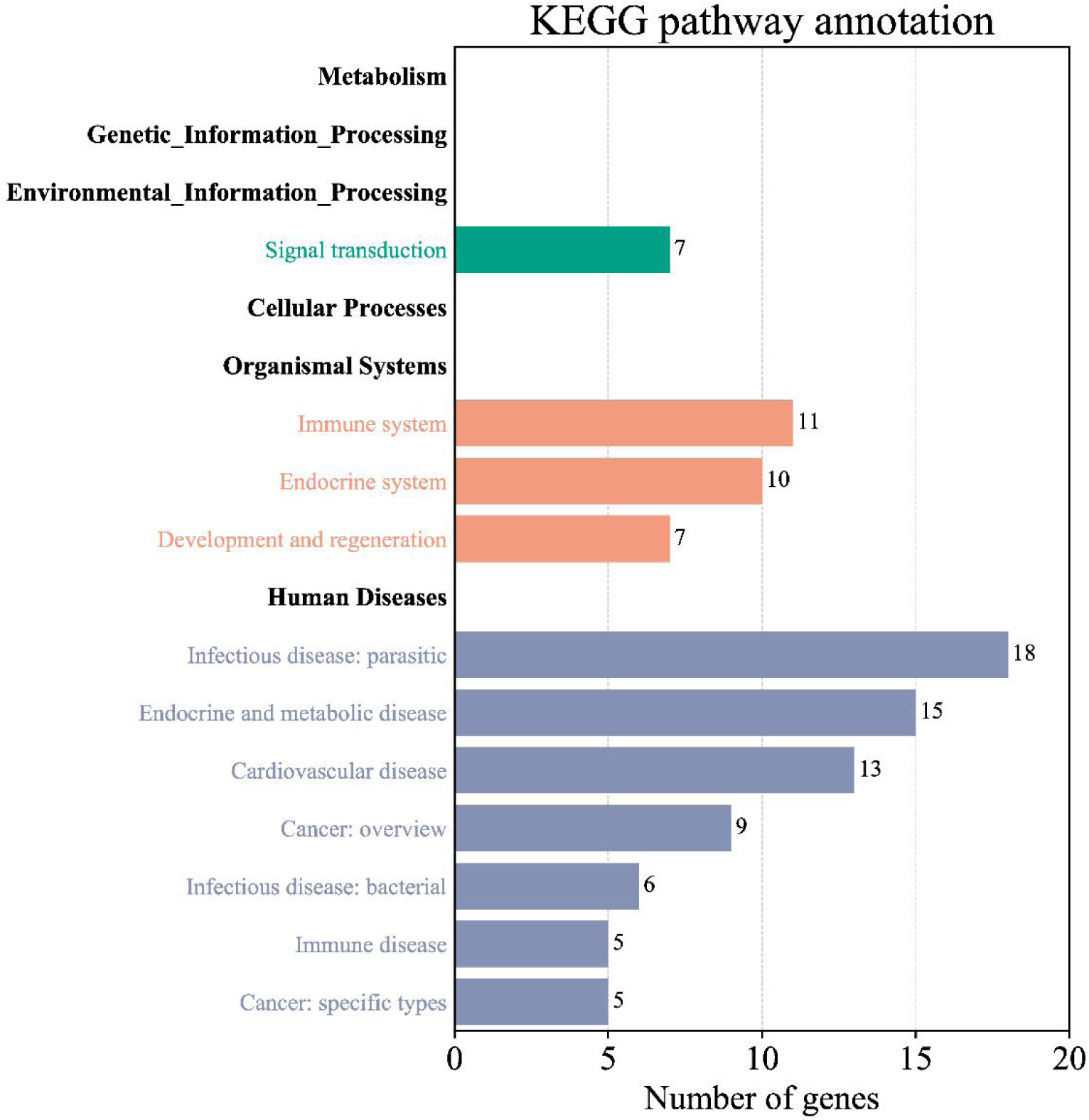
The 10 most significance of KEGG analysis ( 1 and 2 )

**Figure 8:**
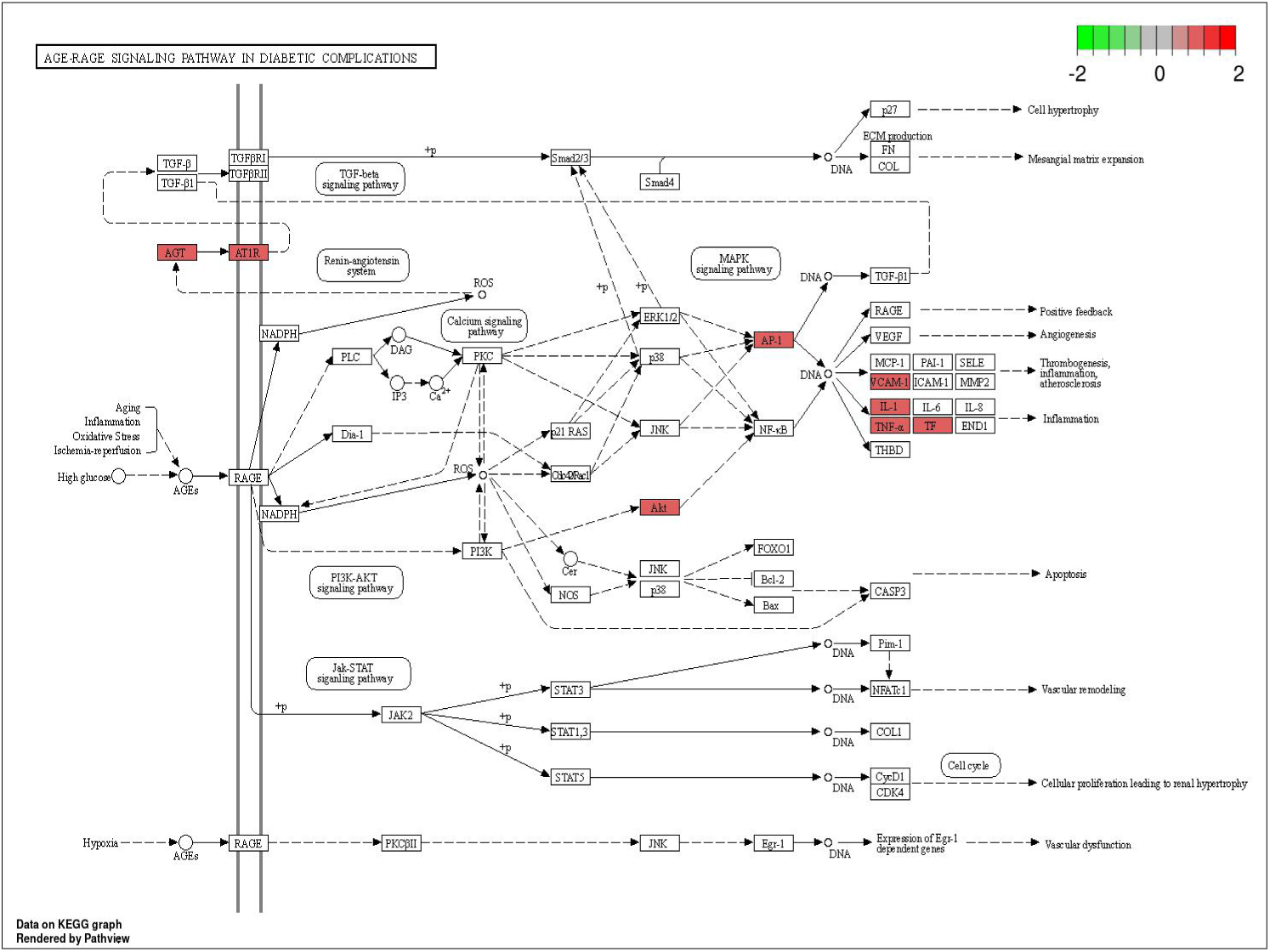
The most prominent pathways and key genes in SSW therapeutic DN targets; □ represents gene products; ○ represents compounds; red represents the key targets of deer antler; → represents intermolecular interactions or relationships.

### 3.5. Component-Target-Pathway

The network relationship of component-target-pathway after the deletion of isolated nodes is shown in Figure 9. As can be seen in the figure, there were a total of 106 nodes, including 5 purple TCM name nodes, 21 green target nodes, 60 blue polygon component nodes, and 20 yellow pathway nodes (Supplement Table 6). The connecting line between the two targets was the edge, which contained 326 edges in total. Each edge represented the interrelation between the medicine-ingredient or ingredient-target or target-pathway. Combined with Figure 1, it can be seen that the treatment of DN with SSW was based on the interaction mode of “multi-component -multi-target - multi-pathway”, rather than only relying on the isolated effect of a single component, a single target, or a single pathway (Figure 9).

**Figure 9:**
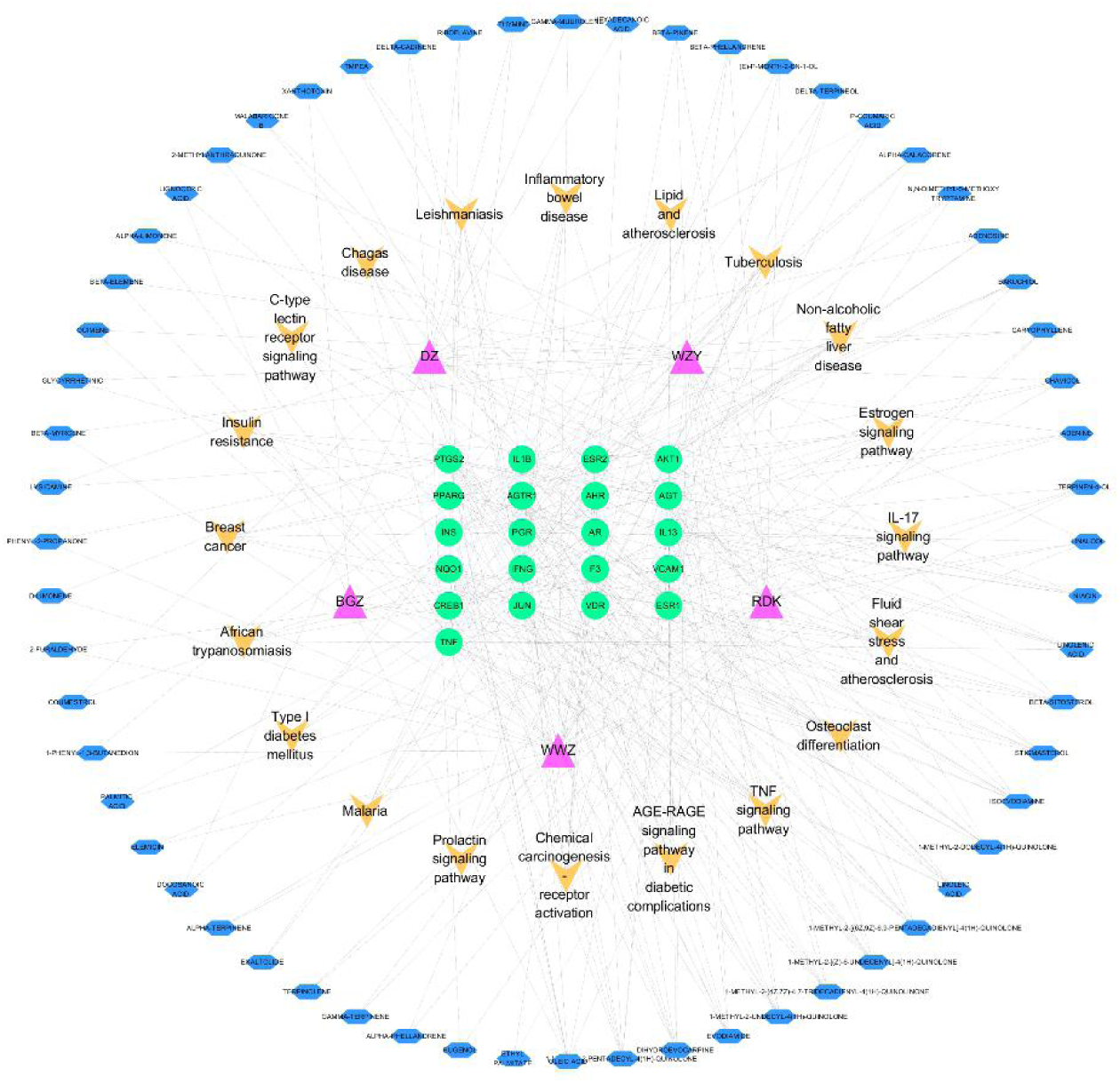
Component-target-pathway network. Blue polygons depict the bioactive components of SSW; green circles denote the targets; yellow circles denote the signaling pathways of DN.

### 3.6. Molecule Docking

To verify the reliability of the component-hub gene interaction, five kinds of hubs were selected as targets for molecular docking. The binding energy of the interaction was shown in the heat map Supplement Table 7. If binding energy <0 kcal·mol-1, it indicated that the ligand molecule could spontaneously bind the receptor protein, while if binding energy < -5.0 kcal·mol-1, it indicated that the ligand molecule had ideal binding force, and the value greater than 7.0 indicated strong binding activity^[30]^. As shown in Figure 10, the binding energy of all binding pairs in this study was < -5kcal·mol-1, indicating that the major bioactive compounds and hub genes had good binding affinity. JUN, PTGS2, TNF, IL-1B, and AKT1 are widely regarded as the classical therapeutic targets of DN. The drug-target binding affinity and optimal docking locations between hub genes and five representative compounds are shown in Figure 11 and Supplement Table 8.

**Figure 10:**
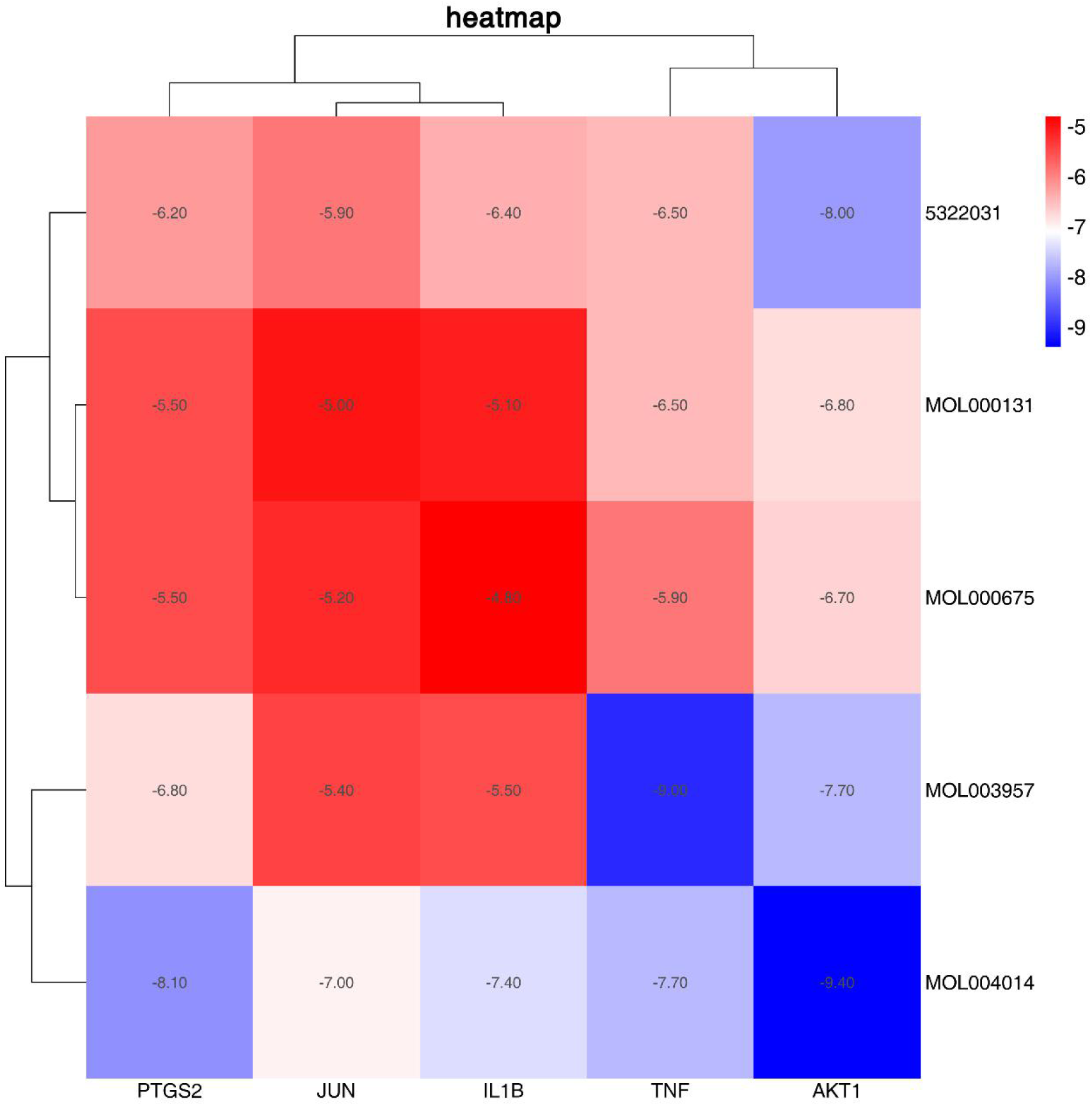
The binding energy values of core compounds of SSW and core targets. 5322031 for DIHYDROEVOCARPINE, MOL000131 for LINOLEIC ACID, MOL000675 for OLEIC ACID, MOL003957 for 1-METHYL-2-PENTADECYL-4(1H)-QUINOLONE, and MOL004014 for EVODIAMIDE.

**Figure 11:**
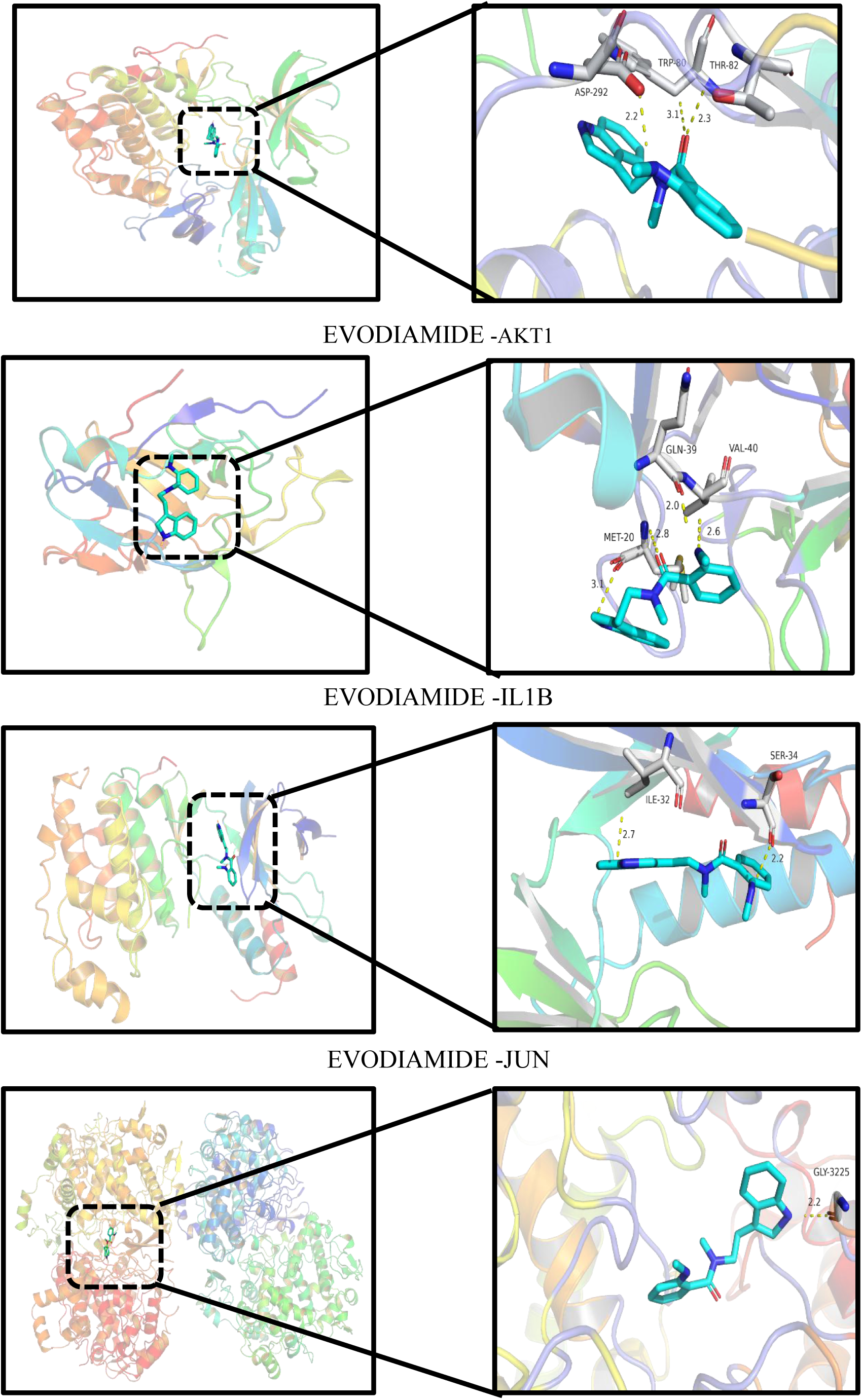
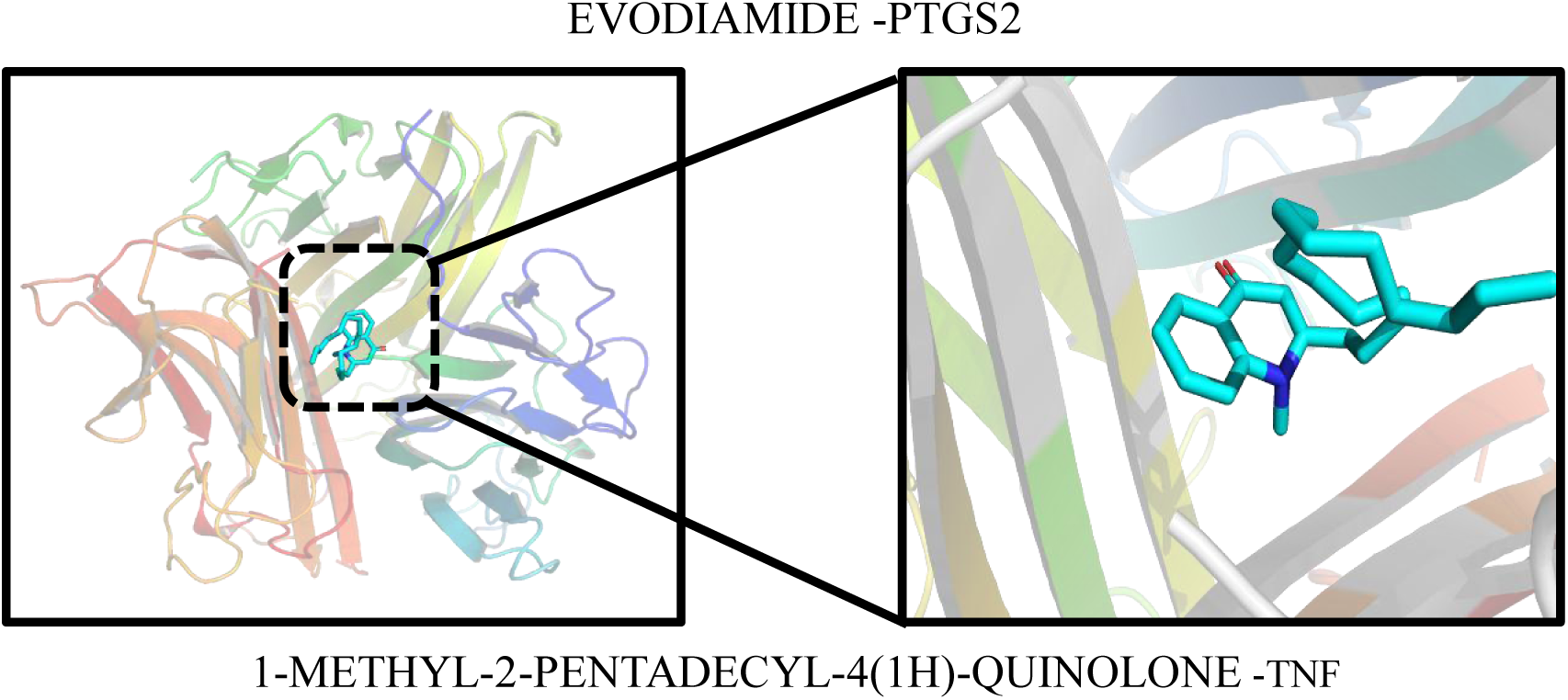
Molecular docking of the optimal conformation for the binding of the 5 hub genes. The dashed blue lines represent hydrogen bonds.

## 4. Discussion

TCM involves complex and diverse compound ingredients, and the single thinking of TCM research can not help fully and systematically understand the role of TCM. Network pharmacology and bioinformatics can systematically study the interaction network of drugs on diseases from a holistic perspective, which can help reveal the mechanism of action of TCM from the molecular level. This coincides with the holistic concept emphasized by TCM. Therefore, in this study, we applied network pharmacology, bioinformatics, and molecular docking to explore the potential mechanism of SSW in the treatment of DN, and successfully screened out 5 main bioactive ingredients, 9 key targets, and 10 important pathways related to SSW.

First of all, 120 effective components corresponding to SSW were screened from TCMSP and BATMAN-TCM online platforms, and 542 corresponding TCM targets were obtained through targeted fishing. By intersecting with DN targets, 195 compound targets by which SSW treated DN were obtained.

In the PPI composite target network, the network analysis were performed on 195 composite targets based on MCODE and degree, and then 9 key targets that were closely related to DN were identified, including INS (degree=75), AKT1(degree=73), TNF(degree=73), IL1B(degree=58), PPARG(degree=51), JUN(degree=49), CREB1(degree=40), and PTGS2(degree=39), ESR1(degree=38). Among the first 20 KEGG pathways, AKT1, TNF, and IL1B targets were found in half of the pathways at the same time, indicating that they were the core genes of SSW treatment of DN. AKT1, as a serine/threonine kinase, can activate the regulatory transduction system in various renal cells, such as podocyocytes, mesangial cells, and renal tubular epithelial cells^[31]^. Besides, AKT1 is a vital regulator of the PI3K/AKT pathway and TGF-β1 mediated biological processes, and it plays an important role in the regulation of inflammation, oxidative stress, apoptosis, and autophagy in DN^[32]^. TNF and IL1B are the main targets of pro-inflammatory factors, and they jointly participate in the formation of chronic inflammation in DN^[33, 34]^.

KEGG pathway enrichment analysis was conducted on 195 intersection targets, and 113 biological pathways were obtained. These pathways, in addition to DN, were also enriched in other diseases possibly because different diseases involved the same target genes. On the one hand, it indicated that SSW might have a potential therapeutic effect on these diseases or that the progression of DN was influenced by other diseases. On the other hand, it indicated that SSW regulated various diseases through multi-target and multi-pathway. This was also a controversial point of network pharmacology. Therefore, the pathways that were closely related to DN were selected in our study, which mainly involved immune response, inflammatory response, regulatory cell function, and oxidative stress. Combined with P value, enrichment gene number and enrichment fraction, and relevant signaling pathways such as AGE-RAGE signaling pathway in diabetic complications, TNF signaling pathway, IL-17 signaling pathway, and fluid shear stress and atherosclerosis were identified as the key pathways in which SSW played a role. Key genes such as AKT1, TNF, IL1B, and JUN were enriched in AGE-RAGE signaling pathway in diabetic complications, and this pathway had the highest enrichment degree and the lowest P value and was closely related to other pathways. Hence, SSW played a key role in delaying the progression of DN. Studies have pointed out that a long-term hyperglycemic environment can increase the oxygen consumption of renal tissue, thus causing hypoxia, inducing the HIF-1-mediated metabolic pathway of renal hypoxia, and even accelerating renal tubule interstitial fibrosis^[35, 36]^. According to related reports, under high glucose conditions, non-enzymatic glycation of proteins and lipids can be induced to produce advanced glycation end products (AGEs), and oxidative stress and inflammation can be induced through activation of AGE receptor (RAGE), aggravating the progression of DN^[37]^. Furthermore, RAGE activation induces the activation of different intracellular signaling pathways, such as PI3K-Akt, MAPK-ERK, and NF-κB^[36]^. Therefore, blocking AGE-RAGE axis formation has become a new treatment strategy for DN at present^[36]^. Studies have revealed that activation of the immune system and the development of chronic inflammation are closely related to the pathogenesis of DN, and the pro-inflammatory environment of diabetes can accelerate the progression of DN^[35]^. Both TNF signaling pathway and the IL-17 signaling pathway, as classical inflammatory pathways, can promote the adhesion and aggregation of inflammatory factors, induce an inflammatory response to participate in microvascular lesions, and ultimately impair renal function^[38]^. Meanwhile, damaged kidney cells can trigger the immune system response, activate a variety of immune pathways such as Toll-like receptor (TLR), NOD-like receptor (NLR), and C-type lectin receptor (CLR), further promote the synthesis of inflammatory cytokines, trigger the chronic inflammatory response of kidney, and lead to the occurrence and progression of DN^[38]^. Fluid shear stress and atherosclerosis pathway includes nuclear factor kappa-B (NF-κB), MAPK and other major signaling pathways, which have been confirmed to be closely related to T2DM^[37]^. Chronic systemic inflammation and activation of innate immunity are regarded as one of the key factors promoting the occurrence and development of T2DM and its complications^[39]^. DN is related to systemic and local renal immune inflammation, in which key inflammatory cells, inflammatory factors and pathways play an important role, including macrophages, nuclear transcription factor-KB (NF-kB), and transcription activator (JAK/STAT) pathway^[40, 41]^. NF-κB, a classical signaling pathway related to inflammation, plays a vital role in improving insulin resistance and inhibiting the apoptosis of islet beta cells and the complications of diabetes. Moreover, AGE-RAGE pathway can activate NF-κB, thus resulting in tissue damage and inflammation, which is a significant mechanism for the occurrence of T2DM and its complications^[42, 43]^. Therefore, the targeted treatment of AGE-RAGE signaling pathway in diabetic complications, TNF signaling pathway, IL-17 signaling pathway and fluid shear stress and atherosclerosis pathways are beneficial in the treatment of DN.

In the composite network diagram of TCM-component-target-pathway, OLEIC ACID, LINOLEIC ACID, 1-METHYL-2-PENTADECYL-4(1H)-QUINOLONE, DIHYDROEVOCARPINE, and EVODIAMIDE are identified as the core components of SSW for the treatment of DN. JUN, PTGS2, TNF, IL-1B and AKT1 are considered to be the first five hub genes of DN that degree value-based SM act, which may play a key role in the therapeutic effect of SSW. In addition, the 5 genes are basically consistent with the core targets obtained in PPI. It has been reported that OLEIC ACID has a certain effect on the softening of blood vessels and plays a vital role in the metabolism of humans and animals. However, OLEIC ACID synthesized by the human body can not meet the needs and should be taken from food, so taking the oil with high LINOLEIC ACID content is healthful^[44]^. As an unsaturated fatty acid, LINOLEIC ACID can reduce blood cholesterol and prevent atherosclerosis^[45]^. Studies have revealed that LINOLEIC ACID is considered as a potential drug for alleviating diabetic complications^[46]^. In addition, clinical experiments have proved that food high in LINOLEIC ACID can reduce plasma lipids in the DN model and inhibit eicosanoic acid synthesis through platelets and renal cortex, thus improving DN model^[46]^. Some studies also point out that LINOLEIC ACID has anti-inflammatory and anti-fibrosis effects in diabetic nephropathy, mediated by inhibiting the increased expression of MCP-1 and ICAM-1 in diabetic conditions^[47]^. In a word, the treatment of DN with SSW is achieved through a multi-component, multi-target, and multi-pathway approach. The relationship between these active compounds and DN deserves further studies.

Finally, based on the results of molecular docking simulation and calculation of binding energy, the binding energy of 25 core component-hub gene pairs was lower than 0, indicating that each pair had a good binding affinity. Evodiide-akt1 docking pair showed the tightest binding (-9.4), followed by 1-METHYL-2-PENTADECYL-4(1H)-QUINOLONE-TNF (-9) and 1-METHYL-2-PENTADECYL-4(1H) -QUINOLONE-PTGS2 (-8.1). The results of molecular docking verified that the core components had good binding characteristics with the hub genes.

## 5. Conclusion

In summary, based on the analysis through network pharmacology and bioinformatics, the results revealed that SSW has a potential molecular biological mechanism on DN through 120 active components and 113 pathways. AGE-RAGE signaling pathway in diabetic complications may be the key to this mechanism. Subsequent molecular docking verification shows that the binding of DN core bioactive compounds with hub genes presents satisfactory results, which further reveal that SSW plays an important role in the treatment of DN. Therefore, this study preliminarily reflects the multi-component, multi-target, and multi-pathway characteristics of SSW, and provides new evidence and a theoretical basis for the research and development of new drugs.

## Limitations

Further pharmacological experimental support is needed for follow-up studies.

## Ethical statement

As a study based on network pharmacology, ethical approval and informed consent of patients are not applicable.

## Author contributions

**Conceptualization:** Yuanyuan DENG, Mianzhi ZHANG.

**Data curation:** Yuanyuan DENG, Sai ZHANG, Yu MA.

**Funding acquisition:** Mianzhi ZHANG.

**Writing – original draft:** Yuanyuan DENG.

**Writing – review & editing:** Mianzhi ZHANG.

## Acknowledgments

We acknowledge the GEO database for providing its platforms and contributors for uploading their meaningful datasets. Meanwhile, we thank Dr. Mianzhi ZHANG, Chief Physician, Dongfang Hospital, Beijing University of Chinese Medicine, for his financial assistance with this study.

## Abbreviations

SSW: Sishenwan
DN: Diabetic Nephropathy
TCMSP: Traditional Chinese Medicine Systems Pharmacology
DL: Drug-likeness
OB: Oral bioavailability
GO: Gene ontology
BP: Biological process
ALB: Albumin
KEGG: Kyoto Encyclopedia of Genes and Genomes
PTGS2: Prostaglandin-endoperoxide synthase 2
CREB1: cAMP-response element binding protein
ESR1: Estrogen receptor 1
TNF: tumor necrosis factor
IL1B: Interleukin-1β
INS: insulin
AKT1: AKT serine/threonine kinase 1
PPARG: Peroxisome proliferator-activated receptor gamma
CC: Cellular component
MF: Molecular Function
TCM: Traditional Chinese medicine

## Supplement materials

**S1 Table. 120 active compounds and 542 corresponding targets of SSW.**

**(PDF)**

**S2 Table. DN-related targets from public databases.**

**(PDF)**

**S3 Table. 195 overlapping targets between SSW and DN.**

**(PDF)**

**S4 Table. Topology information for PPI networks.**

**(PDF)**

**S5 Table. KEGG pathways analysis results.**

**(PDF)**

**S7 Table. Component-Target-Pathway specific information.**

**(PDF)**

**S8 Table. Molecular docking of 5 bioactive compounds and top 5 targets.**

**(DOCX)**

**(PDF)**

## Data availablity statement

Data are available within the GEO database (GSE30528, GSE1009, GSE96804 and GSE47183). STRING repository (https://string-db.org/) and DAVID online database (https://David.ncifcrf.gov/summary.jsp) are the two databases used to process the data and are all online sites that the reader can access and use directly to process the data, as described in the “Methods and Materials” section of the article.

## References

[1] Alicic R Z, Rooney M T, Tuttle K R. Diabetic Kidney Disease: Challenges, Progress, and Possibilities [J]. Clinical Journal of the American Society of Nephrology, 2017: CJN.11491116.

[2] Lacava V, Pellicanò V, Ferrajolo C, et al. Novel avenues for treating diabetic nephropathy: new investigational drugs [J]. Expert Opin Investig Drugs, 2017, 26 (4): 445.

[3] Wang J, Wong Y K, Liao F. What has traditional Chinese medicine delivered for modern medicine? [J]. Expert Rev Mol Med, 2018, 20: e4.

[4] Lu Z, Zhong Y, Liu W, et al. The Efficacy and Mechanism of Chinese Herbal Medicine on Diabetic Kidney Disease [J]. J Diabetes Res, 2019, 2019: 2697672.

[5] Yang X, Hu C, Wang S, et al. Clinical efficacy and safety of Chinese herbal medicine for the treatment of patients with early diabetic nephropathy: A protocol for systematic review and meta-analysis [J]. Medicine (Baltimore), 2020, 99 (29): e20678.

[6] Wang L, Decong M A, Sun L. The Origin and New Exploration of SiShen Pills [J]. Western Journal of Traditional Chinese Medicine, 2015:

[7] Siegel K R, Ali M K, Zhou X, et al. Cost-effectiveness of Interventions to Manage Diabetes: Has the Evidence Changed Since 2008? [J]. Diabetes Care, 2020, 43 (7): 1557.

[8] Tu X, Ye X, Xie C, et al. Combination Therapy with Chinese Medicine and ACEI/ARB for the Management of Diabetic Nephropathy: The Promise in Research Fragments [J]. Curr Vasc Pharmacol, 2015, 13 (4): 526.

[9] DeFronzo R A, Reeves W B, Awad A S. Pathophysiology of diabetic kidney disease: impact of SGLT2 inhibitors [J]. Nat Rev Nephrol, 2021, 17 (5): 319.

[10] Wang Y, Yang H, Chen L, et al. Network-based modeling of herb combinations in traditional Chinese medicine [J]. Brief Bioinform, 2021, 22 (5):

[11] Gauthier J, Vincent A T, Charette S J, et al. A brief history of bioinformatics [J]. Brief Bioinform, 2019, 20 (6): 1981.

[12] Ru J, Li P, Wang J, et al. TCMSP: a database of systems pharmacology for drug discovery from herbal medicines [J]. J Cheminform, 2014, 6: 13.

[13] Xu X, Zhang W, Huang C, et al. A novel chemometric method for the prediction of human oral bioavailability [J]. Int J Mol Sci, 2012, 13 (6): 6964.

[14] Daina A, Michielin O, Zoete V. SwissTargetPrediction: updated data and new features for efficient prediction of protein targets of small molecules [J]. Nucleic Acids Res, 2019, 47 (W1): W357.

[15] Szklarczyk D, Santos A, von Mering C, et al. STITCH 5: augmenting protein-chemical interaction networks with tissue and affinity data [J]. Nucleic Acids Res, 2016, 44 (D1): D380.

[16] Mendez D, Gaulton A, Bento A P, et al. ChEMBL: towards direct deposition of bioassay data [J]. Nucleic Acids Res, 2019, 47 (D1): D930.

[17] UniProt: the universal protein knowledgebase in 2021 [J]. Nucleic Acids Res, 2021, 49 (D1): D480.

[18] Otasek D, Morris J H, Bouças J, et al. Cytoscape Automation: empowering workflow-based network analysis [J]. Genome Biol, 2019, 20 (1): 185.

[19] Cline M S, Smoot M, Cerami E, et al. Integration of biological networks and gene expression data using Cytoscape [J]. Nat Protoc, 2007, 2 (10): 2366.

[20] Piñero J, Ramírez-Anguita J M, Saüch-Pitarch J, et al. The DisGeNET knowledge platform for disease genomics: 2019 update [J]. Nucleic Acids Res, 2020, 48 (D1): D845.

[21] Barshir R, Fishilevich S, Iny-Stein T, et al. GeneCaRNA: A Comprehensive Gene-centric Database of Human Non-coding RNAs in the GeneCards Suite [J]. J Mol Biol, 2021, 433 (11): 166913.

[22] Amberger J S, Bocchini C A, Schiettecatte F, et al. OMIM.org: Online Mendelian Inheritance in Man (OMIM®), an online catalog of human genes and genetic disorders [J]. Nucleic Acids Res, 2015, 43 (Database issue): D789.

[23] Davis A P, Grondin C J, Johnson R J, et al. Comparative Toxicogenomics Database (CTD): update 2021 [J]. Nucleic Acids Res, 2021, 49 (D1): D1138.

[24] Clough E, Barrett T. The Gene Expression Omnibus Database [J]. Methods Mol Biol, 2016, 1418: 93.

[25] Ritchie M E, Phipson B, Wu D, et al. limma powers differential expression analyses for RNA-sequencing and microarray studies [J]. Nucleic Acids Res, 2015, 43 (7): e47.

[26] Kolde R, Laur S, Adler P, et al. Robust rank aggregation for gene list integration and meta-analysis [J]. Bioinformatics, 2012, 28 (4): 573.

[27] Szklarczyk D, Gable A L, Nastou K C, et al. The STRING database in 2021: customizable protein-protein networks, and functional characterization of user-uploaded gene/measurement sets [J]. Nucleic Acids Res, 2021, 49 (D1): D605.

[28] Kim S, Chen J, Cheng T, et al. PubChem in 2021: new data content and improved web interfaces [J]. Nucleic Acids Res, 2021, 49 (D1): D1388.

[29] Burley S K, Bhikadiya C, Bi C, et al. RCSB Protein Data bank: Tools for visualizing and understanding biological macromolecules in 3D [J]. Protein Sci, 2022, 31 (12): e4482.

[30] Zhuang L, Zhai Y-y, Yao W-f, et al. The mechanism study of protecting kidney of Erzhi Pill based on network pharmacology [J]. Acta Pharmaceutica Sinica, 2019: 877.

[31] Heljić M, Brazil D P. Protein kinase B/Akt regulation in diabetic kidney disease [J]. Front Biosci (Schol Ed), 2011, 3 (1): 98.

[32] Khokhar M, Roy D, Modi A, et al. Perspectives on the role of PTEN in diabetic nephropathy: an update [J]. Crit Rev Clin Lab Sci, 2020, 57 (7): 470.

[33] Navarro J F, Mora-Fernández C. The role of TNF-α in diabetic nephropathy: Pathogenic and therapeutic implications [J]. Cytokine & Growth Factor Reviews, 2006, 17 (6): 441.

[34] Buraczynska M, Ksiazek K, Wacinski P, et al. Interleukin-1β Gene (IL1B) Polymorphism and Risk of Developing Diabetic Nephropathy [J]. Immunol Invest, 2019, 48 (6): 577.

[35] Jin Q, Hao X F, Xie L K, et al. A Network Pharmacology to Explore the Mechanism of Astragalus Membranaceus in the Treatment of Diabetic Retinopathy [J]. Evid Based Complement Alternat Med, 2020, 2020: 8878569.

[36] Sanajou D, Ghorbani Haghjo A, Argani H, et al. AGE-RAGE axis blockade in diabetic nephropathy: Current status and future directions [J]. Eur J Pharmacol, 2018, 833: 158.

[37] Wang H, Huang X, Xu P, et al. Apolipoprotein C3 aggravates diabetic nephropathy in type 1 diabetes by activating the renal TLR2/NF-κB pathway [J]. Metabolism, 2021, 119: 154740.

[38] Lampropoulou I T, Stangou Μ, Sarafidis P, et al. TNF-α pathway and T-cell immunity are activated early during the development of diabetic nephropathy in Type II Diabetes Mellitus [J]. Clin Immunol, 2020, 215: 108423.

[39] Macisaac R J, Ekinci E I, Jerums G. Markers of and risk factors for the development and progression of diabetic kidney disease [J]. Am J Kidney Dis, 2014, 63 (2 Suppl 2): S39.

[40] García-García, Patricia M. Inflammation in diabetic kidney disease [J]. World Journal of Diabetes, 2014, 5 (4): 431.

[41] Pérez-Morales R E, Del Pino M D, Valdivielso J M, et al. Inflammation in Diabetic Kidney Disease [J]. Nephron, 2019, 143 (1): 12.

[42] Ke G, Chen X, Liao R, et al. Receptor activator of NF-κB mediates podocyte injury in diabetic nephropathy [J]. Kidney Int, 2021, 100 (2): 377.

[43] Yang H, Xie T, Li D, et al. Tim-3 aggravates podocyte injury in diabetic nephropathy by promoting macrophage activation via the NF-κB/TNF-α pathway [J]. Mol Metab, 2019, 23: 24.

[44] Ying L, Jia Z, Liu S, et al. Combined losartan and nitro-oleic acid remarkably improves diabetic nephropathy in mice [J]. Am J Physiol Renal Physiol, 2013, 305 (11): F1555.

[45] Vergroesen A J. Dietary fat and cardiovascular disease: possible modes of action of linoleic acid [J]. Proc Nutr Soc, 1972, 31 (3): 323.

[46] Barcelli U O, Weiss M, Beach D, et al. High linoleic acid diets ameliorate diabetic nephropathy in rats [J]. American Journal of Kidney Diseases, 1990, 16 (3): 244.

[47] Kim D H, Yoo T H, Lee S H, et al. Gamma Linolenic Acid Exerts Anti-Inflammatory and Anti-Fibrotic Effects in Diabetic Nephropathy [J]. Yonsei Medical Journal, 2012, 53 (6):

